# BRD2 upregulation as a pan-cancer adaptive resistance mechanism to BET inhibition

**DOI:** 10.1101/2025.08.30.673282

**Authors:** Suyakarn Archasappawat, Juliette Jacques, EunJung Lee, Chang-il Hwang

## Abstract

Bromodomain and extraterminal motif (BET) inhibitors, such as JQ1, are promising cancer therapeutics that target epigenetic regulators, particularly BRD4. However, resistance to BET inhibitors (BETi) limits their clinical utility, necessitating a better understanding of adaptive mechanisms. We identified BRD2 upregulation as a conserved response to BET inhibition across multiple cancer types and hypothesized that BRD2 compensates for BRD4 loss, sustaining essential transcriptional programs upon treatment. Consistent with this, BRD2 knockdown sensitized cancer cells to BETi *in vitro*, and combining BRD2 depletion and JQ1 treatment significantly impaired tumor growth *in vivo*. At the chromatin level, BRD2 and BRD4 ChIP-seq analysis of pancreatic cancer cells showed consistent BRD4 loss from chromatin after JQ1 treatment, while BRD2 displacement differed by sensitivity. Resistant cells maintained higher BRD2 occupancy than sensitive cells, suggesting a link between BRD2 retention and drug response. Mechanistically, NFYA mediates BRD2 upregulation as NFYA depletion attenuated BRD2 upregulation upon BETi treatment. Collectively, our findings establish BRD2 as a critical mediator of pan-cancer adaptive resistance to BETi and identify NFYA as a novel transcriptional regulator of this process. Co-targeting BRD2 or its regulatory network offers a rational strategy to enhance the durability and efficacy of BET-based therapies.

## Introduction

Cancer progression can be driven by epigenetic reprogramming, which has recently been recognized as a fundamental hallmark of cancer (1, 2). Epigenetic alterations enable cancer cells to activate oncogenic programs, silence tumor suppressors and adapt to microenvironmental stress, thereby promoting cancer progression and metastasis (3). Importantly, while conferring plasticity and survival, these changes render cancer cells dependent on aberrant epigenetic regulation to sustain the malignant phenotype—vulnerabilities that can be therapeutically exploited. Central to this process is the molecular machinery that regulates epigenetic states, coordinated by three major classes of proteins: writers, readers, and erasers. Writers deposit modifications such as DNA methylation or histone acetylation, erasers remove them, and readers interpret these marks to regulate downstream transcriptional activity. Together, these mechanisms dynamically shape transcriptional programs, often hijacked during carcinogenesis.

Among these regulators, the BET (bromodomain and extra-terminal domain) family has drawn significant attention as a group of critical epigenetic readers. This family comprises four members—BRD2, BRD3, BRD4, and the testis-specific BRDT—all sharing a key structural feature: two tandem N-terminal bromodomains that recognize and bind acetylated lysine residues on histones to anchor transcriptional machinery at active chromatin (4, 5). Of these, BRD4 is the most extensively studied and plays a critical role at enhancer and super-enhancers (SE), large regulatory clusters that drive the expression of key oncogenes, such as *MYC* (6). In addition to tethering to acetylated chromatin, BRD4 also recruits and scaffolds the Mediator complex and the P-TEFb kinase to release paused RNA polymerase II, thereby sustaining high-level transcription of target genes (7). Collectively, the actions of BRD4 support self-reinforcing transcriptional programs driven by inherently unstable oncogenic *MYC* transcripts (3). Because direct pharmacologic inhibition of MYC has proven difficult, BRD4 represents an attractive therapeutic vulnerability. Small-molecule BET inhibitors (BETi) such as JQ1 competitively bind to the bromodomains—most potently of BRD4—displacing them from chromatin and dismantling SE-driven transcription (8). This results in rapid *MYC* downregulation and potent antitumor effects in preclinical models, with notable success in MYC-addicted hematologic malignancies and an increasing exploration across diverse cancer types (9–11).

Despite these promising preclinical outcomes, clinical responses to BETi have been modest and short-lived, with resistance posing as a major barrier for translation (12–14). To date, most mechanistic insights into BETi resistance have come from models of acquired resistance following prolonged drug exposure, in which cancer cells undergo extensive transcriptional and epigenetic rewiring. Such models may not reflect the clinically relevant dynamics of adaptive resistance, defined by early and reversible changes that enable cancer cells to tolerate BET inhibition. A deeper understanding of the early adaptive responses to BET inhibition could reveal vulnerabilities that precede the emergence of acquired resistance and guide the development of more effective upfront or combination therapies.

In this study, we uncovered upregulation of BRD2—a BRD4 paralog—as an adaptive response to BET inhibition across multiple cancer types. Functionally, BRD2 knockdown (KD) sensitizes tumors to the BETi JQ1, enhancing therapeutic efficacy in both *in vitro* and *in vivo* models. Mechanistically, this upregulation is driven, at least in part, by increased expression of the transcription factor NFYA. Notably, our results indicate that the extent of the initial BETi response could help predict the benefit of combining BRD2 KD with JQ1. While all BETi-treated cells benefit from the combination, the most substantial enhancements are observed in initially resistant populations. These findings provide a rationale for incorporating BRD2 targeting into combination strategies to enhance BETi potency and prolong their clinical benefit across diverse cancers.

## Materials and methods

### Cell lines and cell culture conditions

A diverse panel of mouse and human cancer cell lines was used in this study. All cells were maintained at 37°C in a humidified incubator with 5% CO_2_ under standard conditions appropriate for each cell type. The base culture media included Dulbecco’s Modified Eagle Medium (DMEM; Corning, 10-013-CV), RPMI-1640 (Gibco, 11875119), Iscove’s Modified Dulbecco’s Medium (IMDM; Gibco, 12440053), McCoy’s 5A Modified Medium (Gibco, 16600082), Eagle’s Minimum Essential Medium (EMEM; ATCC, 30-2003), and Advanced DMEM/F-12 (Gibco, 12634028). Unless otherwise specified, culture media were supplemented with 10% fetal bovine serum (FBS; GenClone, 25-550H) and 1% penicillin-streptomycin (P/S; Gibco, 15140-122). Triple-negative breast cancer (TNBC) cell lines were additionally supplemented with 10 mM HEPES (Gibco, 15630080), 1 µg/ml hydrocortisone, and 5 µg/ml insulin (Sigma-Aldrich, I0516-5ML). Cell line information, including species, tumor type, tissue origin, specific culture conditions, and corresponding sources and catalog numbers, is listed in the Table S1.

### Drug compounds and treatment conditions

All compounds were purchased from the suppliers listed in Table S2 and dissolved in dimethyl sulfoxide (DMSO; Sigma-Aldrich, 472301-500ML) prior to use as stock solutions. Unless otherwise specified, treatments were performed at final concentration of 1 μM for 24 hours in mouse samples and 10 μM for 24 hours in human samples. For control conditions, cells were treated with DMSO at the same final concentration used to dissolve drug compounds.

### Cell viability assay

Mouse cancer cells and A549, U251, and U2OS cells were seeded at 1,500 cells/well, while all other human cancer cells were seeded at 3,000 cells/well in 50 µl of complete medium into 96-well plates. Twenty-four hours after cell plating, drugs were serially diluted in complete medium and 50 µl of the diluted compounds was added to each well. Cells were incubated for an additional 72 hours before media were carefully removed. To assess cell viability, 100 µl of diluted AlamarBlue solution prepared by mixing 200 µl of resazurin (Fisher Scientific, AC418900010) in 50 ml of 1× PBS was added to each well. Plates were incubated at 37°C with 5% CO_2_ for 3–4 hours. Absorbance at 570 nm was measured using a plate reader (SpectraMax iD5 plate reader, Molecular Devices). Relative cell viability was calculated by normalizing the absorbance of drug-treated wells to DMSO-treated controls. Dose–response curves were generated using nonlinear regression (curve fitting) in GraphPad Prism version 10.6.0 (796).

### Colony formation assay

One thousand cells were seeded per well in 6-well plates. After 1–3 days, cells were treated with JQ1 (50–100 nM) for 7–10 days, with fresh drug-containing medium replaced every 2–3 days. At the endpoint, cells were fixed with 100% methanol (15 minutes, room temperature) and stained with 0.1% (w/v) crystal violet (Spectrum chemical, TCI-C0428-100G) in 20% methanol. Plates were rinsed under running tap water until excess stain was no longer visible, air-dried completely, and scanned (HP LaserJet Pro). Colony area and staining intensity were quantified in ImageJ using the ColonyArea plugin.

### shRNA construct generation via pLKO.1 cloning

For the generation of shRNA constructs, the pLKO.1-puro vector (Addgene #8453) was used. A total of 6 µg of plasmid DNA was digested with 1 µl EcoRI-HF (NEB, R3101T), 1 µl AgeI-HF (NEB, R3552S), and 5 µl rCutSmart Buffer (NEB, B6004S) in a 50 µl reaction and incubated at 37°C for 3 hours. To prevent vector self-ligation, 1 µl Quick CIP (NEB, M0525S) was added and incubated for an additional hour at 37°C. The digested vector was purified using the PureLink PCR purification kit (Invitrogen, K310002). For oligonucleotide preparation, 1 µl of forward and reverse primers (each at 100 µM) were combined with 1 µl 10× T4 DNA ligase buffer (NEB, B0202S), 0.5 µl T4 Polynucleotide Kinase (NEB, M0201S), and 6.5 µl nuclease-free water. The reaction was incubated at 37°C for 30 minutes, then heated to 95 °C for 5 minutes, followed by a gradual ramp down to 25°C at 5 °C/minute for annealing. The annealed primers were diluted 1:200 before ligation. Ligation reactions were performed by mixing 50 ng of digested pLKO.1 vector, 1 µl diluted annealed oligos, 1 µl 10× T4 ligase buffer (NEB, B0202S), 1 µl T4 DNA ligase (NEB, M0202S), and water to a final volume of 10 µl. The mixture was incubated at room temperature for 30 minutes. For transformation, 3 µl of the ligation mix was added to 30 µl XL10-Gold ultracompetent cells (Agilent, 50-125-094). After 5 minutes on ice, cells were heat-shocked at 42°C for 90 seconds, returned to ice for another 5 minutes, and plated on LB agar (Fisher Scientific, BP1427-500) supplemented with 1× ampicillin (RPI, C802G69) using 4.5 mm sterile rattler plating beads (Zymo Research, S1001). Plates were incubated overnight at 37°C (∼16 hours). At least 6 individual colonies were miniprepped using Zyppy Plasmid Miniprep Kit (Zymo Research, D4019). Plasmid inserts were verified by Sanger sequencing (GENEWIZ) using primers targeting the U6 promoter. Resulting sequences were aligned to reference in Benchling to confirm correct oligo insertion. Information about the short hairpin RNA (shRNA) constructs used in this study, including the target gene, sequence ID, and oligonucleotide sequences, is summarized in the Table S3.

### Lentiviral production and transduction

Lentiviral particles were produced by transfecting HEK293T cells using a standard third-generation packaging system. Briefly, 5 × 10⁶ HEK293T cells were seeded in 10 cm culture dishes. After 18–24 hours, the medium was replaced with DMEM supplemented with 10% FBS (without P/S). Cells were then transfected with 10 µg of transfer plasmid (e.g., shRNA constructs), 8 µg of psPAX2 (Addgene #12260), and 2 µg of pMD2.G (Addgene #12259), using X-tremeGENE 360 transfection reagent (Roche, 8724156001) at a 1:3 DNA-to-reagent ratio, following the manufacturer’s instructions. After 48–72 hours, the viral supernatant was collected, centrifuged at 1,100 rpm for 5 minutes to remove cell debris, and filtered through a 0.45 µm PES filter (Cytiva, 28143-312). The resulting virus-containing media were used to transduce target cells in the presence of polybrene (MilliporeSigma, C788D57) at a final concentration of 1 µg/ml. Twenty-four hours after transduction, cells were selected with 2 µg/ml puromycin (Thermo Scientific, 53-79-2) for at least 72 hours. Successful gene knockdown was subsequently validated by Western blot, confirming effective transduction and target suppression.

### siNFYA transfection and JQ1 treatment

NFYA-specific siRNA (Santa Cruz Biotechnology, sc-29947) and non-silencing control siRNA (Santa Cruz Biotechnology, sc-37007) were purchased from Santa Cruz Biotechnology. Cells were transfected with 40 nM siRNA using X-tremeGENE 360 reagent (Roche, 8724156001) following the manufacturer’s protocol. After 48 hours, transfected cells were treated with JQ1 (1 µM for 24 hours), and NFYA knockdown efficiency was assessed by Western blot.

### Western blotting

Cells at 80–90% confluency were scraped on ice using a cell lifter (Biologix, 70-2180) into lysis buffer (50 mM Tris, pH 7.5; 150 mM NaCl; 5 mM EDTA; 1% Triton X-100) supplemented with protease inhibitor cocktail (Sigma-Aldrich, SIAL-11836170001) and phosphatase inhibitor cocktail (Roche, 04906837001). Lysates were incubated on ice for 30 minutes and then clarified by centrifugation at 14,000 rpm for 10 minutes at 4°C. Supernatants were transferred to fresh tubes, and protein concentrations were measured using the DC Protein Assay Kit (Bio-Rad; Reagents A, B, S: 5000113, 5000114, 5000115). A total of 20 μg of protein was mixed with 4× LDS sample buffer (Invitrogen, B0007) and 10× NuPAGE Sample Reducing Agent (Invitrogen, NP0009), adjusted to a final volume of 20 μl, and heated at 70°C for 10 minutes. Samples were loaded onto 4–12% Bis-Tris NuPAGE gels (Invitrogen, NP0321) and electrophoresed using a Mini Gel Tank (Thermo Fisher, A25977) in MES SDS Running Buffer (Invitrogen, NP0002) at 120 V for 90 minutes. Proteins were transferred to 0.45 μm PVDF membranes (Millipore, IPVH00010) at 350 mA for 150 minutes at 4°C. Membranes were blocked in 5% skim milk in Tris-buffered saline (Bio-Rad, 1706435) with 0.1% Tween-20 (Fisher Scientific, BP337-500) (TBST) for 1 hour at room temperature. They were then incubated overnight at 4°C with primary antibodies (see Table S4) diluted in 5% skim milk in TBST under gentle agitation (∼70 rpm) (ONILAB, SK-O180-S). Following incubation, membranes were washed three times for 10 minutes each with 1× TBST on a rocker and then incubated with HRP-conjugated secondary antibodies (see Table S4) in the blocking buffer for 1 hour at room temperature. Membranes were subsequently washed three more times for 10 minutes with TBST. Signal was detected using SuperSignal West Femto Maximum Sensitivity Substrate (Thermo Scientific, 34094) and visualized using the Amersham Imager 600 (GE Healthcare Life Sciences). Knockdown efficiency was quantified in ImageJ using the Gel Analysis tool by measuring band intensities (area under the curve). Target protein levels were normalized to the loading control and expressed relative to shScr for each cell line.

### Animal studies and treatment regimen

Female athymic nude (*Nu/Nu*) mice (6 weeks old) were obtained from the Jackson Laboratory (Strain #:002019) and housed under specific pathogen-free conditions in the Genome and Biomedical Sciences Facility at the University of California, Davis. Mice were maintained with controlled ambient temperature and a standard light–dark cycle. All procedures were approved by the University of California Davis Institutional Animal Care and Use Committee (Protocol #24016). JQ1 (MedChemExpress, HY-13030) was prepared by dissolving the drug in DMSO (Sigma-Aldrich, 472301-500ML), then diluting in vehicle composed of 95% of a 20% SBE-β-CD solution (MedChemExpress, HY-17031) in saline (Moltox, C790D04) to a final concentration of 5 mg/ml. For subcutaneous tumor engraftment, cells were trypsinized, passed through a 40 μm cell strainer (Biologix, 15-1040), and resuspended in 50 μl of Cultrex Reduced Growth Factor Basement Membrane Extract (Bio-Techne, 3533-010-02). Each mouse received 0.5 × 10⁶ cells: shScramble (shScr) control cells in the left flank and shBrd2 cells in the right flank. Three days post-engraftment, mice were randomized into two treatment groups (n = 10 per group) and administered either vehicle or JQ1 intraperitoneally at 50 mg/kg, once a day, five days per week. Tumor size was measured using Vernier calipers three times per week, and tumor volume was calculated as: (π×length×width²)/6. At the end of the treatment period, mice were euthanized, and tumors were harvested for histological analysis.

### Histology and immunohistochemistry

Harvested tumors were fixed in 10% neutral buffered formalin (Fisher Scientific, 22-110-873) and replaced with 70% ethanol (Thomas Scientific, C993Y81) on the following day. Formalin-fixed paraffin-embedded sections with 5 µm thickness were prepared for immunohistochemistry (IHC), which was performed at the Center for Genomic Pathology, University of California, Davis. For Ki67 staining, antigen retrieval was performed for 45 minutes with citrate buffer at pH 6.0 in a Decloaking Chamber (Biocare Medical) at 125°C and 15 psi. Slides were blocked with normal goat serum then incubated with rabbit polyclonal Ki67 (NeoMarkers, RB1510P, 1:1000 dilution) overnight at room temperature in a humidified chamber, followed by biotinylated goat anti-rabbit secondary antibody (1:1000 dilution). The Vectastain^®^ ABC elite kit and diaminobenzidine peroxidase substrate kit (Vector Laboratories) were used for amplification and visualization of signal, respectively Ki67 staining was quantified in ImageJ using the IHC Profiler plugin (15). Nuclei classified as high positive or positive by the plugin were counted as Ki67-positive.

### RNA extraction

Cells were lysed in 1 ml of TRIzol reagent (Thermo Fisher Scientific, 15596026) at room temperature for 5 minutes with gentle agitation, then 200 µl of chloroform (Reagents, C755N59) was added. Total RNA was purified using the PureLink RNA Mini Kit (Invitrogen, 12183018A) according to the manufacturer’s instructions. RNA concentration and purity were assessed using a NanoDrop ND-1000 spectrophotometer (Thermo Scientific) prior to downstream applications.

### Reverse transcription quantitative PCR (RT-qPCR)

Total RNA was extracted using TRIzol reagent (Invitrogen, 15596026). After 5 minutes of incubation at room temperature, 200 µl of chloroform (Reagents, C755N59) was added, and samples were vigorously mixed until homogeneous, followed by 5 minutes on ice. Samples were centrifuged at 14,000 rpm for 15 minutes at 4°C, and the aqueous phase was transferred to a new DNase/RNase-free tube. RNA was further purified using the PureLink RNA Mini Kit (Invitrogen, 12183018A), including on-column DNase treatment (PureLink DNase Set, Invitrogen, 12185010). RNA was eluted in nuclease-free water, and concentration and purity were assessed using a NanoDrop ND-1000 spectrophotometer (Thermo Scientific).

cDNA synthesis was performed with the High-Capacity cDNA Reverse Transcription Kit (Applied Biosystems, 4368814) using 1 µg of total RNA per 20 µl reaction. The reaction mix included 3 µl 10× RT buffer, 1.2 µl 25× dNTP mix, 3 µl 10× random primers, 1.5 µl MultiScribe reverse transcriptase, and 6.3 µl nuclease-free water. The thermocycling conditions were: 25°C for 10 minutes, 37°C for 120 minutes, and 85°C for 5 minutes.

qPCR was performed using 1 µl of cDNA with PowerTrack SYBR Green Master Mix (Applied Biosystems, A46109) and gene-specific primers (listed in Table S5) in a total reaction volume of 10 µl. Reactions were run on a LightCycler 480 system (Roche) with the following setting: 95°C for 10 minutes, followed by 40 cycles of 95°C for 15 seconds and 60°C for 60 seconds. Relative mRNA expression levels were normalized to *GAPDH* (human samples) or *Gapdh* (mouse samples) and calculated using the 2^−ΔΔCt^ method.

### Chromatin immunoprecipitation assays (ChIP-seq and ChIP-qPCR)

#### BRD2 and BRD4 ChIP-seq ChIP

was performed using the SimpleChIP^®^ Enzymatic Chromatin IP Kit (Magnetic Beads) (Cell Signaling Technology, 9003) with minor modifications. Briefly, approximately 1 × 10⁷ mT3-2D or mT23-2D cells were dissociated with TrypLE Express (Gibco, 12605028), resuspended in PBS, crosslinked with 1% formaldehyde for 10 minutes at room temperature, and quenched with 125 mM glycine for 5 minutes. Chromatin was digested with MNase (1 µL, 37°C, 20 minutes) and sonicated (Bioruptor, Diagenode) at a high power (3 rounds of 10 cycles, 30 seconds on/30 seconds off) to obtain ∼150–900 bp fragments. A 2% input aliquot (10 µl) was reserved, and the remaining chromatin was incubated overnight at 4°C with constant rotation with 10 µl of BRD2 (Cell Signaling Technology, 5848S) or BRD4 (Bethyl, A301-985A100) antibody per reaction. Bead capture, washes, elution, reverse crosslinking, and DNA purification were performed according to the kit protocol.

#### NFYA ChIP-qPCR

ChIP-qPCR was performed similarly using approximately 4 × 10⁶ HEK293T cells per reaction, except chromatin was digested with 0.5 µl MNase (37°C, 10 minutes) and sonicated for 6 cycles (30 seconds on/30 seconds off at high power). Immunoprecipitations used 1 µl of NFYA antibody (Diagenode, C15310261) or 10 µl IgG control (Cell Signaling Technology, 2729S). qPCR was performed using PowerTrack SYBR Green Master Mix (Applied Biosystems, A46109) with primers in Table S5, following the qPCR conditions described in the RT-qPCR section. Percent input was calculated as 100 × 2^(Ct adjusted input − Ct ChIP)^. ChIP-seq and ChIP-qPCR were generated using a single biological replicate per condition.

### Library preparation and sequencing

Library preparation and sequencing were carried out by Novogene Co., LTD. For RNA-seq, total RNA quality was assessed using 1% agarose gel electrophoresis, NanoDrop spectrophotometry, and an Agilent 2100 Bioanalyzer. Poly(A) mRNA was enriched from total RNA using oligo(dT) magnetic beads, fragmented, and reverse-transcribed to synthesize first-strand cDNA with random primers. Second-strand cDNA synthesis was performed using a dUTP-incorporation protocol to maintain strand specificity. The resulting double-stranded cDNA was end-repaired, A-tailed, and ligated to Illumina adapters, followed by size selection (250–300 bp insert) and PCR amplification. Libraries were quantified, quality-checked, and sequenced on an Illumina NovaSeq X Plus platform to generate paired-end 150 bp reads (PE150). For ChIP-seq, sample quality was assessed prior to library construction. DNA fragments were end-repaired, A-tailed, and ligated to Illumina adapters, followed by size selection, PCR amplification, and quantification. Libraries were sequenced on an Illumina NovaSeq X Plus platform (PE150). Quality control procedures, including assessments of sequencing quality distribution, error rate distribution, and AT/CG base composition, were performed before data delivery.

### Bioinformatic analysis

All analyses were performed on the Galaxy platform unless otherwise stated. Raw reads were quality-trimmed with Trim Galore! to remove adapter sequences and low-quality bases.

#### RNA-seq analysis

For RNA-seq, clean reads were aligned to the *Mus musculus* mm9 reference genome using HISAT2, and gene-level counts were obtained with featureCounts. Normalization and differential expression analysis were performed with DESeq2, using the median-of-ratios method. P-values were adjusted for multiple testing using the Benjamini–Hochberg procedure, and genes with adjusted p < 0.05 were considered significantly differentially expressed. Normalized counts were obtained from DESeq2, annotated using annotatemyIDs, and used for downstream visualization. Volcano plots were generated by plotting the log_2_FC against –log_10_p-adj to highlight significantly upregulated and downregulated genes.

#### ChIP-seq analysis

All ChIP-seq samples were processed using the same analysis pipeline with identical parameters as described below. Trimmed reads were aligned to the *Mus musculu*s mm9 reference genome with Bowtie2, then filtered with SAMtools to retain only properly paired reads with a mapping quality ≥ 30 while excluding mitochondrial reads (chrM). PCR and optical duplicates were removed with Picard MarkDuplicates. To ensure that only high-quality, non-redundant reads were used in downstream analyses, we evaluated ChIP enrichment using standard QC metrics. Across all BRD2/BRD4 ChIP-seq samples, the mean mapped read depth was 17.4 million reads. Enrichment assessed by FRiP averaged 0.33 across IP samples, and IP enrichment relative to input was additionally supported by deepTools plotFingerprint.

Peaks were called with MACS2. Consensus peaks were defined as those present in both biological replicates using bedtools intersect. For each cell line, BRD2/BRD4 co-occupied regions were identified by intersecting BRD2 and BRD4 consensus peak sets. Regions were recentered on peak summits and extended ± 2 kb to create fixed windows for signal extraction.

Signal coverage tracks (BigWig files) were generated from duplicate-removed BAM files using bamCoverage with RPGC normalization (scaled to scaled to 1× genome coverage). Signal changes across ranked regions were visualized with deepTools (plotHeatmap, plotProfile). Per-region signal was extracted using deepTools computeMatrix reference-point (±2,000 bp, 50-bp bins, refPoint = center, skipZeros = true, missingDataAsZero = true). For each region, bins were averaged to obtain a single signal value per replicate and condition (DMSO, JQ1). Using a single ChIP-seq dataset per condition, we used a rank-based, region-level analysis to summarize relative signal changes and interpreted these results as descriptive.

To define BRD2 regions most sensitive to BET inhibition in the sensitive cell line, a per-region change in mT3-2D was calculated as Δ = BRD2_DMSO_ − BRD2_JQ1_, where larger positive Δ indicates greater BRD2 loss following JQ1. The top 5% mT3 JQ1-sensitive BRD2 regions (mm9 coordinates) were assigned to candidate genes using GREAT with the default basal-plus-extension model (proximal: 5 kb upstream and 1 kb downstream of the transcription start site; distal up to 1 Mb), yielding 1,035 associated genes. Functional enrichment for GO Biological Process (GOBP) terms was first assessed using GREAT, and the same gene set was additionally analyzed using ShinyGO GOBP (default mouse genome background; Benjamini–Hochberg FDR correction; q < 0.05) for complementary visualization and reporting. The top 10 enriched terms were reported. For integrative expression analysis, GREAT-associated genes were intersected with the RNA-seq expression matrix (DESeq2-normalized counts), resulting in 1,028 genes that were visualized in Morpheus as heatmaps of row-scaled z-scores (z-score computed per gene across samples).

### Analysis of publicly available datasets

#### RNA-seq analysis of BRD2 response to BET inhibition (JQ1)

RNA-seq expression data from human conventional cancer cell lines were obtained via the GEO2R portal. Differential expression analysis was performed using the limma package as implemented in GEO2R. Log_2_FC and –log_10_p-adj for *BRD2* were extracted to evaluate its response to JQ1 treatment, and these values were visualized as dot plots in R. To visualize differential expression patterns after JQ1 treatment, 3D volcano plots were generated in R (version 4.3.0) using DESeq2 output from GEO2R, with specific focus on *BRD2* and *NFYA*. A summary of the datasets analyzed, including GEO accession numbers, cancer types, and sample details, is provided in the Table S6.

#### RNA-seq analysis of BRD4 knockdown

RNA-seq data from human conventional cancer cell lines were obtained from the GEO2R portal. Normalized counts were used as provided, and BRD2 expression was compared between control and BRD4 knockdown conditions. The datasets used in this analysis are listed in Table S7.

#### ChIP-seq analysis of BRD2, BRD4, and NFYA

BRD2 and BRD4 ChIP-seq data from human breast cancer cell lines (GSE131097) were obtained from GEO (listed in Table S8). Processed peak files (BED) and signal tracks (BigWig) were used directly from the repository. For BRD2 and BRD4 ChIP-seq, binding sites were intersected using bedtools intersect to define co-occupied regions, and signal distribution across these regions was assessed with computeMatrix followed by plotProfile to generate metaprofiles. For NFYA, H3K4me3, RNA Pol II ChIP-seq, browser tracks of each BigWig file from GEO (listed in Table S9) were visualized in IGV (version 2.13.2) at the *BRD2* locus.

### Statistical analysis

All statistical analyses were performed using GraphPad Prism version 10.6.0 (796). Statistical significance between two groups was assessed using a two-tailed, unpaired Student’s t-test. For comparisons involving more than two groups, one-way ANOVA with Tukey’s post hoc test was used. A p < 0.05 was considered statistically significant. Error bars represent the SEM unless otherwise indicated.

## Results

### BET inhibition upregulates BRD2 across cancer types

To identify adaptive transcriptional responses to BET inhibition across cancer types, we first performed a systematic analysis of publicly available transcriptomic data. In total, we curated 51 human RNA-seq datasets following treatment with a small molecule BETi JQ1 from the Gene Expression Omnibus (GEO), published over the past decade (2014–2024), encompassing a wide spectrum of malignancies including prostate, breast, brain, skin, gastrointestinal, lung, pediatric, gynecologic and hematologic cancers (16–64). Across all datasets analyzed, *BRD2* expression was consistently and significantly upregulated upon JQ1 treatment, emerging as a recurrent transcriptional response to BET inhibition **(Fig. 1A)**, while no consistent induction was observed for BRD3 and BRD4. The strongest increase in *BRD2* expression was observed in castration-resistant prostate cancer (CRPC; GSE78213), where JQ1 treatment induced a significant upregulation (log_2_FC = 2.15; p-adj = 0.00), whereas *BRD3* and *BRD4* exhibited only modest changes (log_2_FC = –0.13, p-adj = 0.0931; log_2_FC = 0.15, p-adj = 0.561, respectively) **(Fig. 1B)**. To validate the findings from the publicly available datasets, we selected cell lines from multiple cancer types represented in the GEO analysis and assessed *BRD2* expression after JQ1 treatment. These included CRPC (PC3), hepatocellular carcinoma (HCC; Huh7), triple-negative breast cancer (TNBC; MDA-MB-231 and SUM159), glioblastoma (GBM; U251), neuroblastoma (NB; SK-N-BE2), non-small cell lung cancer (NSCLC; A549), osteosarcoma (OS; U2OS), mouse and human acute myeloid leukemia (AML; RN2 and HL-60). In all cell lines tested, BRD2 protein levels were significantly increased following JQ1 treatment **(Fig. 1C; Fig. S1A)**. Notably, BRD2 expression was generally low in leukemia cell lines that we tested. In human AML (HL-60), BRD2 was undetectable at the protein level under baseline conditions; however, we confirmed its induction at the mRNA level in response to JQ1. Similarly, in mouse AML (RN2), BRD2 protein became detectable only after JQ1 exposure **(Fig. S1B–S1C)**. Collectively, these findings confirmed that BRD2 upregulation emerged as a universal response to BET inhibition, distinguishing it from other BET family members. These results suggest a previously underappreciated role for BRD2 in the context of pharmacological BET blockade and raise the possibility that its upregulation reflects a feedback or compensatory mechanism.

**Fig. 1.**
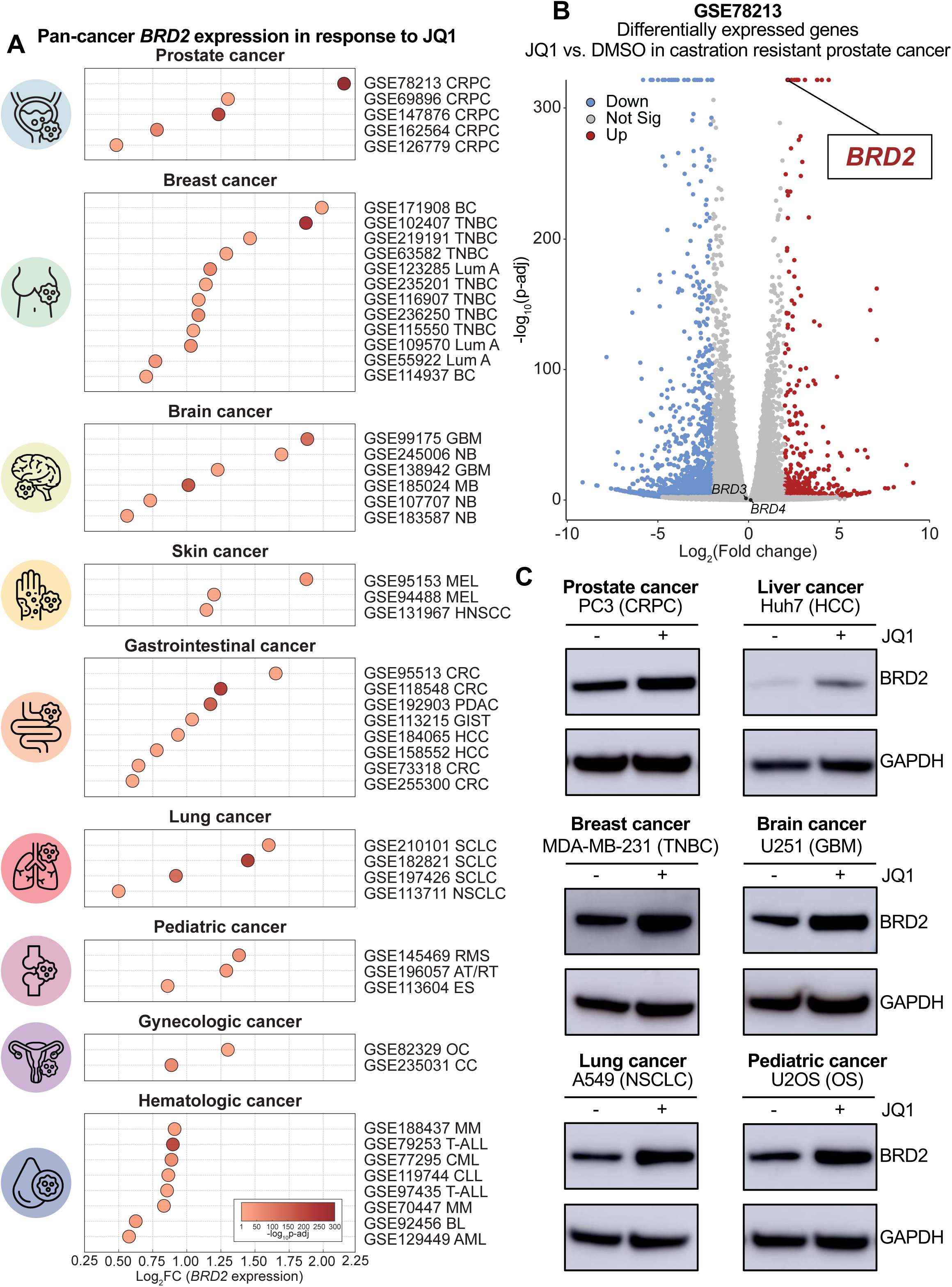
Pan-cancer BRD2 upregulation upon BET inhibition. **(A)** Dot plots showing *BRD2* expression changes (JQ1 vs. control) across 51 curated human RNA-seq datasets from GEO (2014–2024), including gastrointestinal, breast, prostate, skin, brain, lung, pediatric, gynecologic, and hematologic cancers. The x-axis represents log_2_fold change (log_2_FC), and dot size corresponds to statistical significance (–log_10_p-adj). Abbreviations: CRPC, castration-resistant prostate cancer; TNBC, triple-negative breast cancer; Lum A, luminal A; BC, breast cancer; GBM, glioblastoma; NB, neuroblastoma; MEL, melanoma; HNSCC, head and neck squamous cell carcinoma; CRC, colorectal cancer; PDAC, pancreatic ductal adenocarcinoma; GIST, gastrointestinal stromal tumor; HCC, hepatocellular carcinoma; SCLC, small cell lung cancer; NSCLC, non-small cell lung cancer; RMS, rhabdomyosarcoma; AT/RT, atypical teratoid/rhabdoid tumor; ES, Ewing sarcoma; OC, ovarian cancer; CC, cervical cancer; MM, multiple myeloma; T-ALL, T-cell acute lymphoblastic leukemia; CML, chronic myeloid leukemia; BL, Burkitt lymphoma; AML, acute myeloid leukemia. **(B)** Volcano plot of differentially expressed genes in CRPC (GSE78213), highlighting robust *BRD2* upregulation upon JQ1 treatment. *BRD3* and *BRD4* remained unaffected. **(C)** Western blot validation showing BRD2 protein levels in multiple representative human cell lines from prostate (PC3), liver (Huh7), breast (MDA-MB-231), brain (U251), lung (A549), and pediatric (U2OS) cancers with and without JQ1.

### Pharmacological BET inhibition, not BRD4 depletion, induces BRD2 upregulation in pancreatic cancer

Previously, we reported that BRCA2 deficiency sensitized pancreatic ductal adenocarcinoma (PDAC) cells to BETi (65), in contrast to the general resistance observed in most PDAC cells to BETi. To further investigate the underlying mechanisms of *BRD2* upregulation upon BET inhibition, we used PDAC as a model system. In the publicly available dataset GSE192903, which profiles PANC-1 cells treated with JQ1, BRD2 was significantly upregulated (log_2_FC = 1.18, p-adj = 4.87 × 10^−^¹⁸⁵) **(Fig. 2A)**. To validate this, we treated 3 murine KPC pancreatic cancer cell lines (mT3-2D, mT19-2D, and mT23-2D) with JQ1 and conducted RNA-seq. Principal component analysis (PCA) revealed distinct transcriptomic shifts upon the treatment, indicating that JQ1 broadly altered the transcriptional landscape **(Fig. S2A)**. In line with their transcriptional activation role, JQ1 treatment led to widespread gene downregulation, with 1,839 genes significantly decreased. Although *Brd3* and *Brd4* expression remained unchanged, 353 genes were significantly upregulated. Notably, *Brd2* was among the most significantly induced genes (log_2_FC = 2.25, p-adj = 7.16 × 10^−^²⁵) **(Fig. 2B)**. This finding was further validated by RT-qPCR and Western blot in both murine primary and metastatic PDAC cells, along with a panel of human PDAC cell lines, confirming that BRD2 upregulation is a robust and reproducible transcriptional response to BET inhibition across species and disease stages **(Fig. 2C–D; Fig. S2B–D)**. Importantly, this effect was not limited to JQ1; treatment with other small-molecule BETi, including birabresib, molibresib, CPI-0610, ZEN-3694, and AZD5153, also elevated BRD2 protein expression **(Fig. 2E; Fig. S2E–F)**. Because BRD4 is a major BET family with well-established role in transcriptional regulation, we next asked whether BRD2 upregulation could be induced by genetic depletion of BRD4. To test this, we knocked down BRD4 using shRNA in the same murine PDAC cell lines. Despite successfully silencing BRD4, BRD2 expression remained unchanged, highlighting that BRD4 KD elicits a different transcriptional response from pharmacological BET inhibition. **(Fig. 2F; Fig. S2G)**. Consistent with this observation, BRD2 was upregulated by JQ1 in various cancer models, but not by genetic depletion of BRD4 **(Fig. S2H)** (47, 55, 66–69). While the transcriptomic profile of shBRD4 cells shifted in a similar direction to that of JQ1-treated cells, the changes were less pronounced, indicating that BRD4 KD only partially phenocopies BET inhibition **(Fig. 2G)**. Together, these findings demonstrate that pharmacological BET inhibition induces BRD2 upregulation as part of a broader transcriptional response that is mechanistically distinct from BRD4 depletion, potentially reflecting an adaptive program to preserve transcriptional output.

**Fig. 2.**
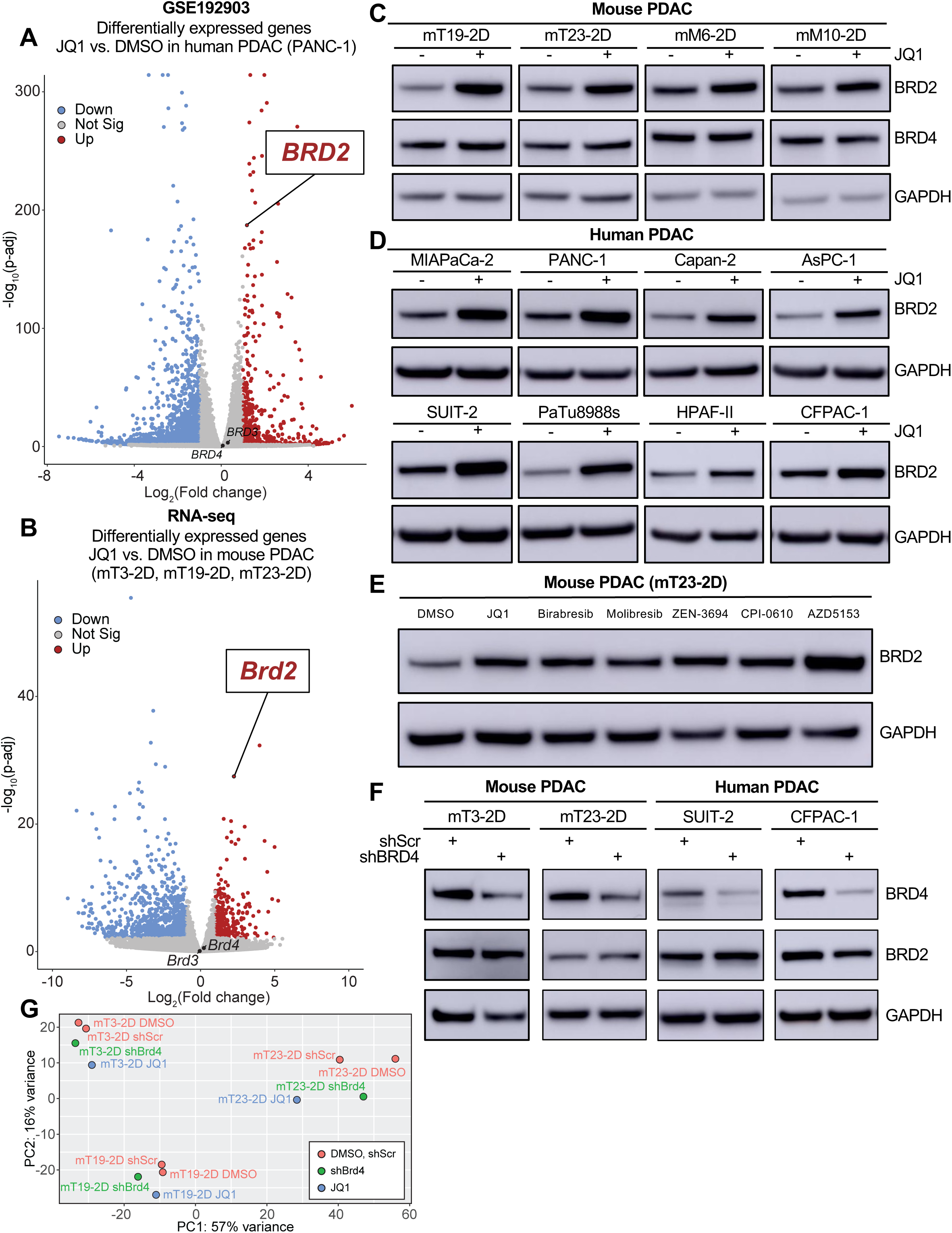
Pharmacological BET inhibition uniquely drives BRD2 upregulation in pancreatic cancer. **(A)** Volcano plot of differentially expressed genes in human PDAC (GSE192903) treated with JQ1, highlighting strong upregulation of *BRD2*, neither *BRD3* nor *BRD4*. **(B)** Volcano plot of RNA-seq data from murine PDAC cells treated with JQ1, showing conserved and significant upregulation of *Brd2*. **(C)** Western blot validation of BRD2 upregulation in murine PDAC cell lines treated with JQ1. **(D)** Western blot analysis in a panel of human PDAC cell lines confirming JQ1-induced BRD2 expression. **(E)** Western blot analysis of BRD2 in murine PDAC cells (mT23-2D) treated with different small-molecule BET inhibitors, showing a consistent increase in BRD2 expression across inhibitors. **(F)** Western blot analysis of BRD2 expression following shRNA-mediated BRD4 knockdown (KD) in murine and human PDAC cell lines, revealing that BRD4 depletion does not mimic JQ1-induced BRD2 upregulation. **(G)** Principal component analysis (PCA) of RNA-seq data from murine PDAC cells under DMSO, JQ1, shScramble (shScr), or shBrd4 conditions, showing distinct transcriptional clustering of JQ1-treated samples compared to BRD4 KD.

### NFYA is one of the potential upstream regulators of BRD2 in response to BET inhibition

BETi-mediated induction of BRD2 prompted us to investigate the upstream factors that might be linked to its regulation. Given that gene induction is often shaped by transcription factors (TF), we sought to nominate TF candidates that might regulate BRD2. We began by querying the COEXPRESdb platform for transcription factors co-expressed with BRD2. Among the top 30 co-expressed genes, Nuclear Transcription Factor Y subunit Alpha (NFYA) was the only TF identified **(Fig. S3A–B)**. Consistent with this, our analysis of transcriptomic data from DepMap across 1,684 human cancer cell lines revealed a positive correlation between *NFYA* and *BRD2* expression (Pearson’s *r* = 0.628, p = 0.00) **(Fig. 3A)**. We next asked whether *NFYA* and *BRD2* exhibit coordinated responses to BET inhibition. Analysis of both publicly available RNA-seq datasets and our mouse PDAC dataset revealed that both *NFYA* and *BRD2* were upregulated under JQ1 treatment **(Fig. 3B; Fig. S3C)**. To evaluate whether NFYA acts as an upstream regulator of *BRD2* transcription, we analyzed public NFYA ChIP-seq datasets and observed NFYA signal near the *BRD2* transcription start site (TSS) within an active promoter region marked by H3K4me3 and RNA Pol II occupancy across multiple cell types **(Fig. 3C; Fig. S3D)**. To test this relationship, we performed siRNA-mediated KD of NFYA in HEK293T and U2OS cells. Depletion of NFYA reduced basal levels of *BRD2* expression and partially reduced BRD2 expression after JQ1 treatment **(Fig. 3D–G; Fig. S3E–F)**. Indeed, NFYA ChIP-qPCR showed the increased NFYA binding at the BRD2 promoter region upon JQ1 treatment relative to IgG control **(Fig. S3D)**, suggesting that NFYA may contribute, at least in part, to *BRD2* transcriptional upregulation in response to BET inhibition.

**Fig. 3.**
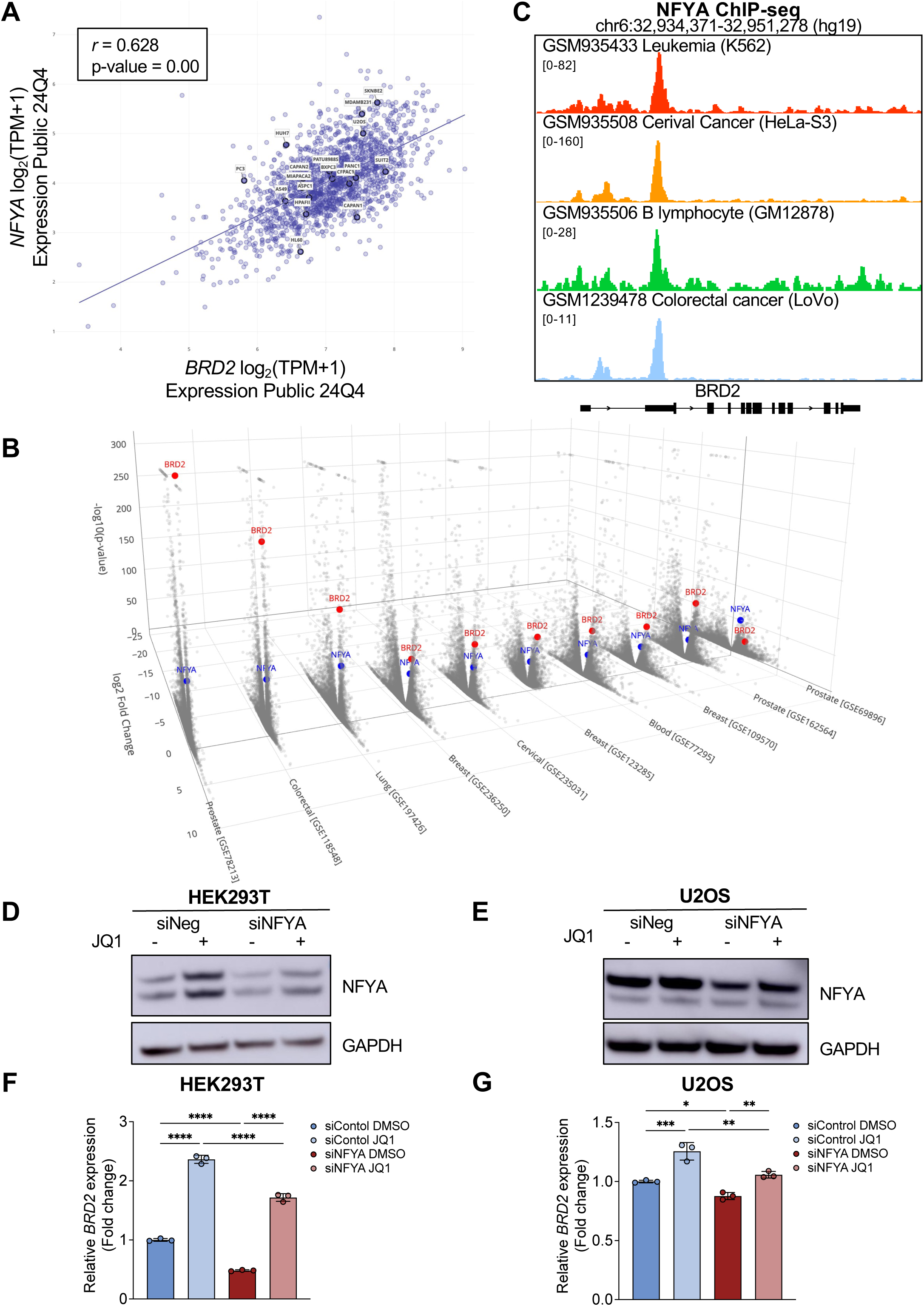
NFYA as a potential candidate transcriptional factor associated with BRD2 induction upon BET inhibition. **(A)** Scatter plot from DepMap analysis showing a positive correlation between *NFYA* and *BRD2* expression across public cancer transcriptomes. **(B)** Representative 3D volcano plot of publicly available RNA-seq datasets treated with JQ1, highlighting co-upregulation of *BRD2* and *NFYA*. **(C)** NFYA ChIP-seq profiles at the *BRD2* locus across multiple cell types. **(D,E)** Western blot analysis of NFYA and BRD2 in HEK293T (D) and U2OS (E) cells transfected with control siRNA (siNeg) or NFYA siRNA (siNFYA) and treated with DMSO or JQ1. **(F,G)** RT-qPCR analysis of *BRD2* expression in HEK293T (F) and U2OS (G) cells under the same conditions, showing that NFYA KD attenuated JQ1-mediated *BRD2* upregulation. Statistical significance was determined by one-way ANOVA (*p < 0.05, **p < 0.01, ***p < 0.001, ****p < 0.0001).

### BRD2 knockdown sensitizes cells to BET inhibition *in vitro*

Given the robust upregulation of BRD2 following BET inhibition, we hypothesized that BRD2 upregulation might confer adaptive resistance to BETi. To test this, we performed shRNA-mediated KD of BRD2 in a panel of cancer cell lines and evaluated their sensitivity to BETi. In mouse PDAC cells, BRD2 KD enhanced sensitivity to 6 different BETi—JQ1, birabresib, molibresib, ZEN-3694, CPI-0610, and AZD5153 **(Fig. 4A–B)**. Similar results were also observed in 3 additional human cancer cell lines—A549 (NSCLC), U251 (GBM), and U2OS (OS)—where BRD2 KD consistently increased JQ1 sensitivity **(Fig. 4C–H)**. Across multiple cell lines, BRD2 depletion sensitized cells to BET inhibition, supporting that BRD2 upregulation contributes to adaptive resistance against BET inhibition **(Fig. S4A–I)**. Moreover, clonogenic assays under chronic treatment mirrored this response, with durable suppression of colony outgrowth over extended BETi exposure **(Fig. S4J–O)**. Interestingly, we observed that the magnitude of sensitization varied depending on the cell’s initial response to BETi. Cells that were more resistant to BET inhibition at baseline (mT23-2D, mM1-2D, mM6-2D, and Capan-2) exhibited a greater reduction in IC_50_ upon BRD2 KD compared to those that were initially sensitive (mT3-2D, CFPAC-1, and BxPC3) **(Fig. 4I–N, Fig. S4A–I)**. In sum, our results revealed a functional role for BRD2 in mediating an acute adaptive response to BETi, suggesting that targeting BRD2 enhances therapeutic efficacy, particularly in BETi-resistant tumors.

**Fig. 4.**
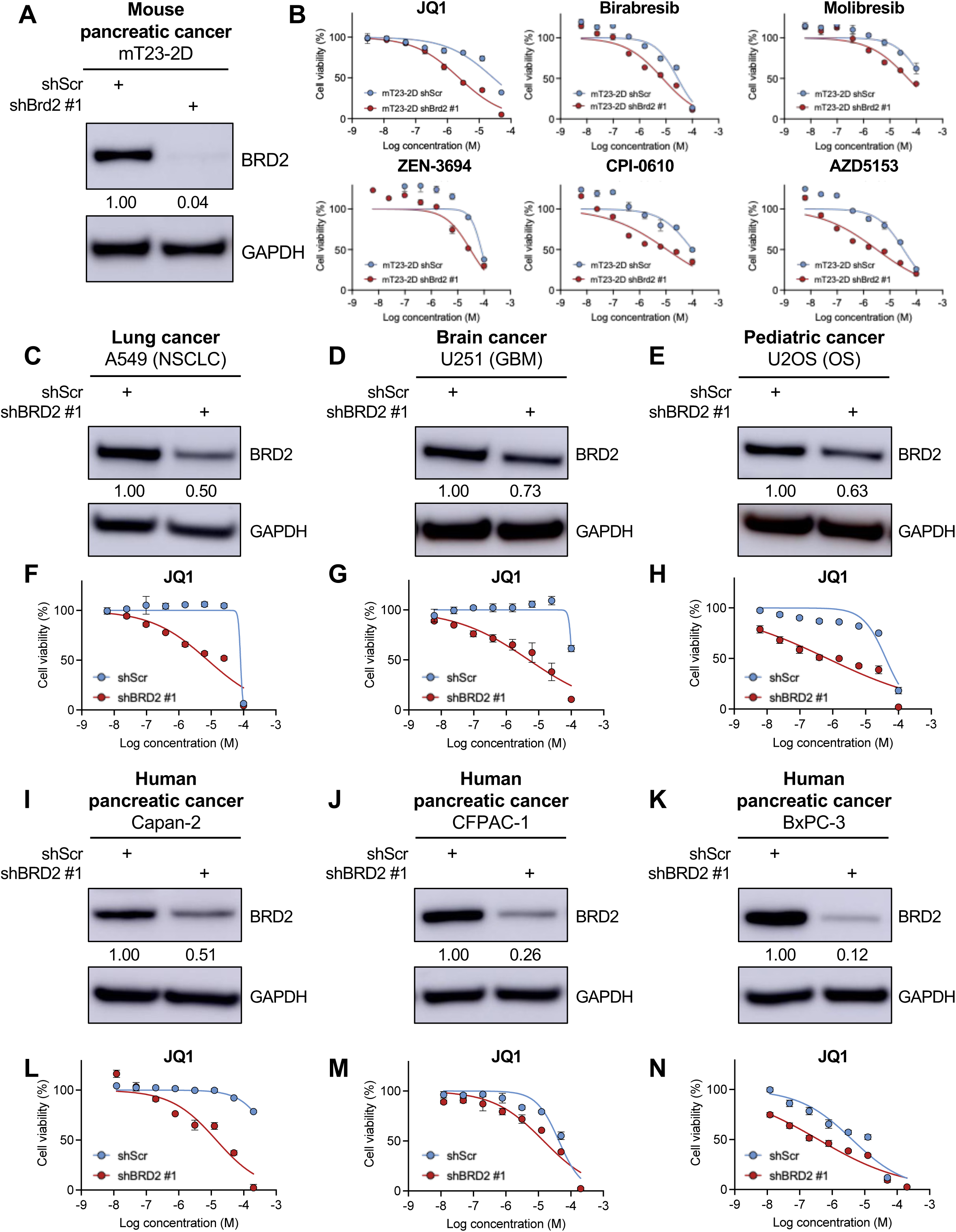
BRD2 depletion sensitizes cancer cells to BET inhibition. **(A)** Immunoblot confirming BRD2 KD (shBRD2 #1) in mT23-2D PDAC cells. Values indicate relative BRD2 protein levels quantified by densitometry, normalized to GAPDH, and expressed relative to shScr. **(B)** Dose–response curves showing reduced cell viability upon treatment with the indicated BETi in BRD2-depleted mT23-2D cells compared to shScr controls. **(C–N)** Immunoblots confirming BRD2 KD (shBRD2 #1) and JQ1 dose–response curves in lung (A549), brain (U251), pediatric (U2OS), and human PDAC (Capan-2, CFPAC-1, and BxPC-3) cell lines following BRD2 KD.

### BRD2 depletion enhances BETi efficacy *in vivo*

To evaluate whether BRD2 contributes to BETi response *in vivo*, we subcutaneously injected mouse pancreatic cancer cells (mT23-2D) harboring control or BRD2-targeting shRNA into immunocompromised athymic *Nu/Nu* mice. Mice were treated intraperitoneally with either vehicle or JQ1 (50 mg/kg) over a three-week period **(Fig. 5A)**. While both JQ1 treatment alone or BRD2 KD alone moderately suppressed tumor growth, combined BRD2 KD and JQ1 treatment resulted in striking tumor growth suppression **(Fig. 5B–D)**. No overt toxicity was observed as body weight remained stable across treatment arms **(Fig. S5)**. Immunohistochemical analysis of Ki67 staining revealed a marked reduction in proliferative cells in BRD2 KD tumors treated with JQ1, which exhibited the lowest percentage of Ki67-positive cells **(Fig. 5E–F)**. Together, these results demonstrate that BRD2 depletion significantly enhances the antitumor efficacy of BET inhibition *in vivo*.

**Fig. 5.**
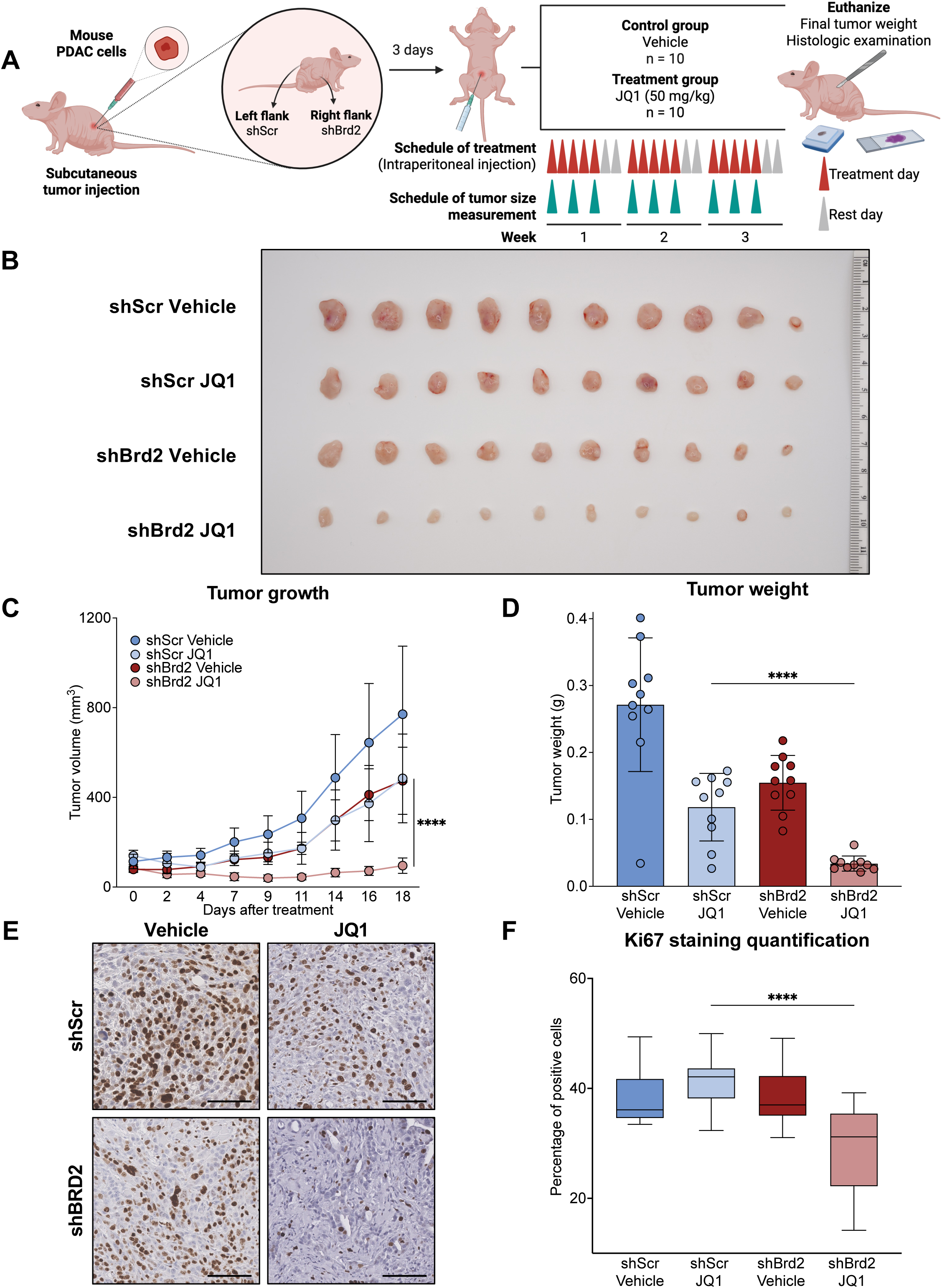
Genetic BRD2 perturbation enhances the antitumor effect of BET inhibition *in vivo*. **(A)** Schematic of *in vivo* experimental design, created with BioRender. **(B)** Representative images of mice and excised tumors from each group after treatment. Scale bar, 1 cm. **(C)** Tumor growth curves over time showing significantly impaired tumor growth in shBrd2 + JQ1 tumors compared to controls. Error bars represent SD. **(D)** Tumor weight at endpoint, showing that shBrd2 + JQ1 tumors had the lowest weights. Statistical significance was determined by two-tailed Student’s t-test (****p < 0.0001). **(E–F)** Representative Ki67 immunohistochemistry and quantification showing significantly reduced proliferative index in shBrd2 + JQ1 tumors compared to controls. Statistical significance was determined using two-tailed Student’s t-test (****p < 0.0001). Scale bar, 100 µm.

### BRD2 persists on chromatin preferentially in BETi-resistant cells

Given our findings that BRD2 is associated with adaptive resistance to BET inhibition both *in vitro* and *in vivo*, we sought to investigate potential underlying mechanisms. We first examined the basal expression levels of BRD4 and BRD2 but did not observe a clear correlation with JQ1 IC_50_ values **(Fig. S6A–B)**, consistent with DepMap data across 394 cell lines (Pearson’s *r* = –0.102 and –0.058 for BRD4 and BRD2, respectively) **(Fig. S6C–D)**. Since BETi displaces BET proteins from acetylated chromatin, we hypothesized that differential BETi responses between BETi-sensitive (mT3-2D) and BETi-resistant (mT23-2D) cells **(Fig. 6A; Fig. S6E)** may reflect differential BRD2 and BRD4 chromatin occupancy after BETi treatment. To address this, we performed BRD2 and BRD4 ChIP-seq in both BETi-sensitive (mT3-2D) and BETi-resistant (mT23-2D) cells to compare their chromatin occupancy under DMSO and JQ1 conditions. In the DMSO condition, we identified common BRD4 (n = 24,810) and BRD2 peaks (n = 23,706) between these two cell lines. Notably, about 70% of BRD4-bound regions (16,941 of 24,810 peaks) were also bound by BRD2 **(Fig. 6B)**. In these overlapping regions, upon JQ1 treatment, we found that BRD4 chromatin occupancy overall decreased in both cell lines to a similar extent. However, BRD2 occupancy was retained to a greater degree in BETi-resistant (mT23-2D) cells than BETi-sensitive (mT3-2D) cells **(Fig. 6C)**. A similar pattern of BRD2 persistence has been observed in TNBC cell line (SUM149) after BET inhibition, suggesting a broader relevance across cancer types **(Fig. S6F–G)**. To further examine whether differential BRD2 displacement affects transcriptional output, we focused on the top 5% of BRD2/BRD4 co-occupied sites showing the greatest loss of BRD2 binding after JQ1 treatment in mT3-2D cells (847 regions), which were comparatively better preserved in JQ1-treated mT23-2D cells. Consistent with this binding pattern, our RNA-seq analysis revealed a more pronounced transcriptional shift for genes associated with these sites in mT3-2D cells, whereas the corresponding transcriptional profiles in mT23-2D cells remained closer to DMSO-treated controls **(Fig. 6D)**. Gene ontology analysis of this gene set showed enrichment for chromatin organization/modification and RNA processing programs, suggesting potential roles of these genes in mediating adaptive resistance of BETi **(Fig. 6E; Fig. S6H)**. Collectively, our data support a model in which, upon BET inhibition, resistant behavior is associated with greater BRD2 chromatin retention and more preserved transcriptional output despite reduced BRD4 occupancy **(Fig. 6F)**. These findings suggest that BRD2 may serve as a compensatory factor when BRD4 is displaced from chromatin and provide a rationale for targeting BRD2 in combination with BETi to improve therapeutic responses.

**Fig. 6.**
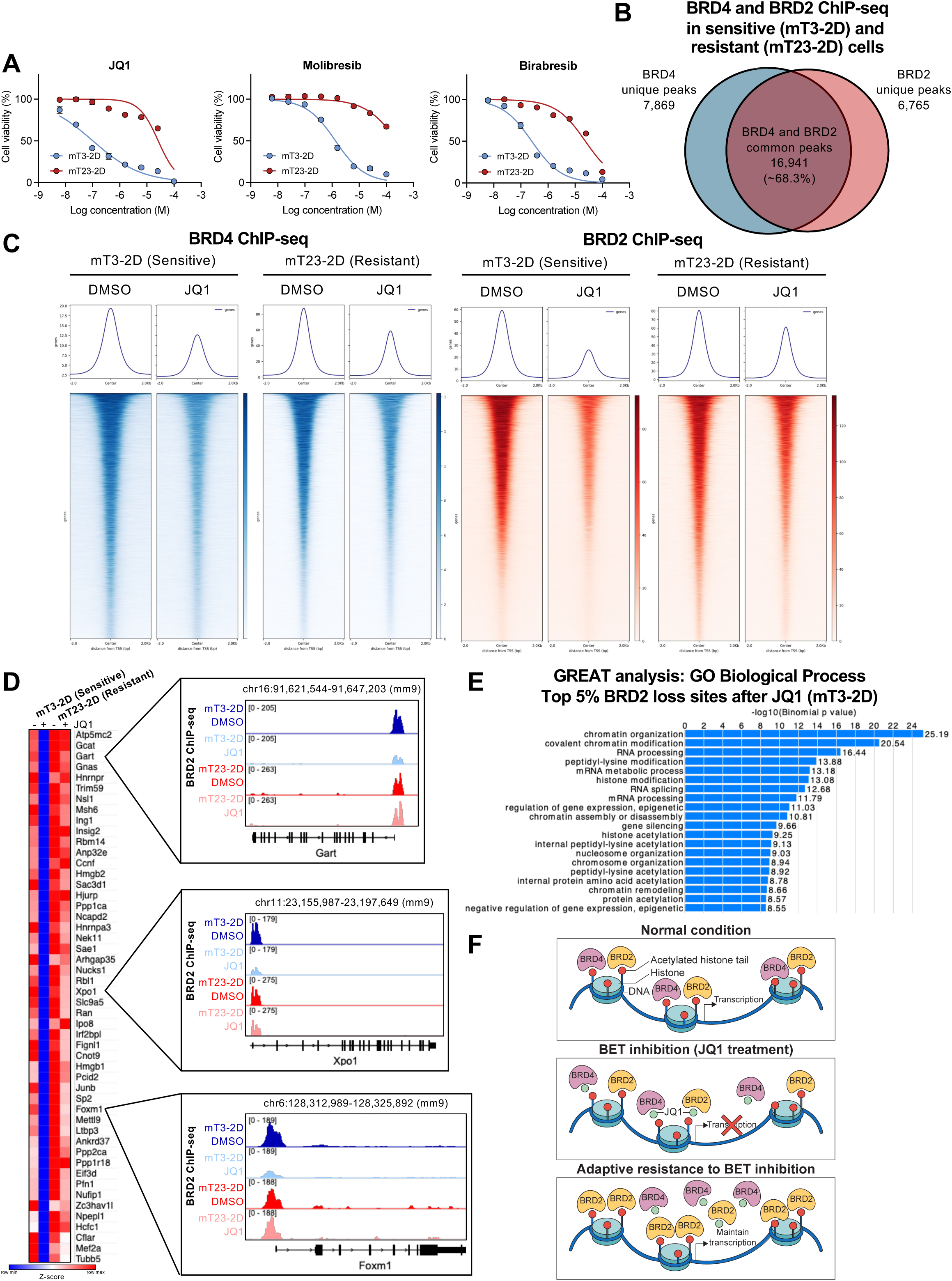
BRD2 chromatin retention may help maintain transcriptional programs in BETi-resistant pancreatic cancer cells. **(A)** Dose–response curves comparing sensitivity to BET inhibitors in BETi-sensitive (mT3-2D) and JQ1-resistant (mT23-2D) PDAC cells. **(B)** Venn diagram showing overlap between BRD4 and BRD2 ChIP-seq peaks across BETi-sensitive (mT3-2D) and BETi-resistant (mT23-2D) cells. **(C)** Metaplots and heatmaps of BRD4 and BRD2 ChIP-seq signal at BRD2/BRD4-overlapping peaks (16,941 regions) in BETi-sensitive (mT3-2D) and BETi-resistant (mT23-2D) cells treated with DMSO or JQ1. **(D)** RNA-seq heatmap (row-scaled z-scores) and BRD2 ChIP-seq browser tracks at representative loci in both BETi-sensitive (mT3-2D) and BETi-resistant (mT23-2D) cells **(E)** GREAT gene ontology enrichment for genes associated with top 5% BRD2-loss sites after JQ1 in mT3-2D cells. **(F)** Proposed schematic model illustrating BRD2 upregulation as an adaptive mechanism to BET inhibition.

## Discussion

BRD4 dependency in many cancers, including leukemia, provided the rationale for developing BETi. The first-generation inhibitor JQ1 demonstrated potent antitumor effects in preclinical models such as NUT carcinoma and MYC-driven leukemia. However, its poor bioavailability, short half-life, and off-target toxicities limited its clinical use, prompting the development of second-generation BETi with improved pharmacokinetics, enhanced specificity, and orally available formulations. Indisputably, despite these advances, cancer cells can activate alternative pathways to circumvent therapy, mitigating its effects and sustaining survival. These adaptive responses not only represent tangible vulnerabilities but, when understood, can also be exploited to prevent resistance and achieve better therapeutic outcomes. Several studies have informally noted increased BRD2 expression following BETi treatment, but it has remained unclear whether this functionally confers pan-cancer adaptive resistance. Recent synthetic lethal screens with BETi in TNBC and paralog-focused CRISPR screens in lung, skin, pancreas, and brain cancers have independently identified BRD2 as a potential resistance factor (70, 71). Yet, the role of BRD2 has not been systematically examined. Here, we show that transcriptional upregulation of BRD2 is a robust adaptive response to BETi across diverse cancer types, and that BRD2 depletion sensitizes cells to BET inhibition both *in vitro* and *in vivo*.

Paralog compensation is an emerging theme in targeted cancer therapy, wherein inhibition of one protein prompts adaptive upregulation of a closely related family member that compensates for its function. For example, EZH2 inhibition in malignant rhabdoid tumors has been shown to induce EZH1 upregulation, and dual EZH1/2 inhibition yields improved therapeutic responses (72). Similarly, inhibition of the histone demethylase LSD1 (KDM1A) in Ewing sarcoma cells resulted in marked induction of its homolog LSD2 (KDM1B), a shift that modulates drug sensitivity (73). Recent large-scale paralog-focused CRISPR screens have uncovered multiple additional examples of this phenomenon, such as the reciprocal buffering between DUSP4 and DUSP6 (71). Here, we present the first comprehensive evidence that BRD2 upregulation is a highly conserved and universal adaptive response to BET inhibition across cancer types, functionally analogous to other paralog compensation events in which BRD2 substitutes for BRD4 to preserve transcriptional programs upon BET inhibition.

Our data suggest that resistance is mediated in part through paralog redundancy, with BRD2 compensating for BRD4 function under BET inhibition. This compensatory role aligns with their shared canonical functions in transcriptional regulation, as evidenced by their largely overlapping genomic occupancy (70). However, BRD2 also carries out distinct functions that may independently and uniquely support adaptive resistance. Upon JQ1 treatment, BET proteins can remain in the nucleus and, through their extra-terminal (ET) domains, may exert bromodomain-independent functions that sustain cell survival. Unlike BRD4, BRD2 is closely associated with chromatin architectural regulation: it co-localizes genome-wide with CTCF, contributes to the maintenance of topologically associating domain (TAD) boundaries, and influences compartmentalization of the accessible genome (74, 75). The C-terminal domain of BRD2 also mediates critical protein-protein interactions, enabling its recruitment to promoters and CTCF-bound sites. The unique roles of BRD2 raise the possibility that its upregulation under BET inhibition may confer survival advantages beyond its role in transcriptional regulation.

Our findings suggest a therapeutic opportunity to enhance BETi efficacy by preventing or counteracting BRD2 upregulation. This could be achieved either through selective BRD2 inhibition in combination with conventional BETi which predominantly target BRD4 or by strategies that broadly degrade BET family members to prevent compensatory mechanisms. PROTAC-based BET degraders, which remove BRD2, BRD3, and BRD4 from chromatin, have recently demonstrated superior antitumor activity compared to earlier generations of BETi (59, 76). Notably, the absence of BRD2 upregulation in PROTAC-based approaches may partially explain their improved efficacy. While additional resistance mechanisms undoubtedly exist, our data highlight BRD2 compensation as a conserved and targetable pathway. As a proof-of-principle, we show that BRD2 depletion sensitizes tumors to BET inhibition. This provides a rationale for integrating BETi with approaches that mitigate BRD2-driven adaptive resistance to achieve more durable clinical responses and expand the therapeutic potential of BET-targeted strategies in cancer therapeutics.

## Supplementary figure legends

**Supplementary Fig. 1.**
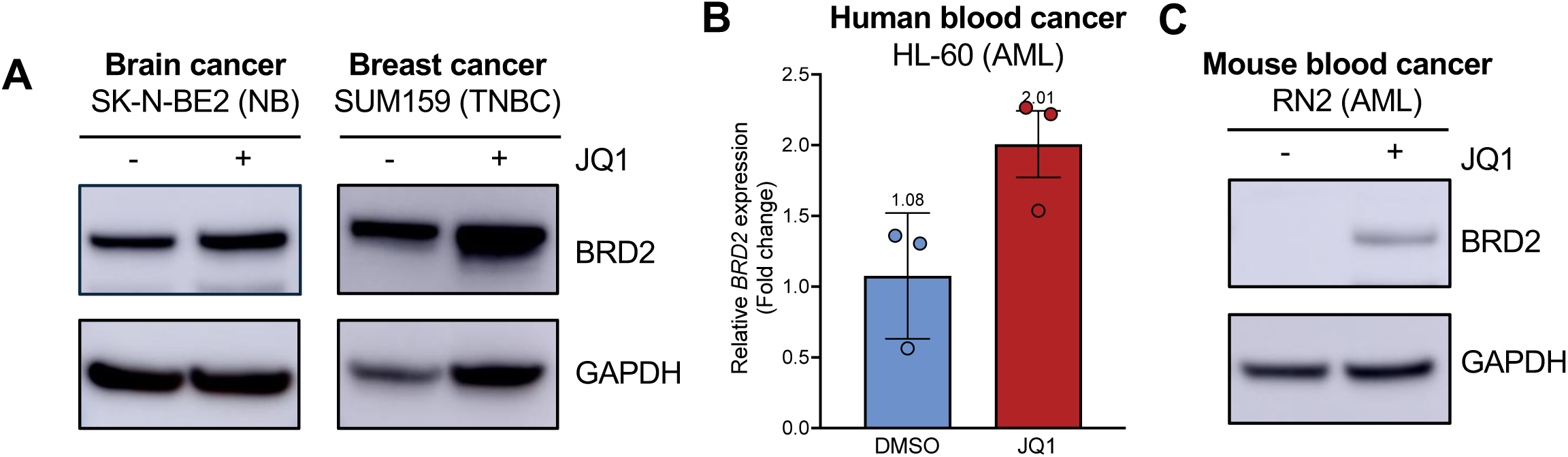
BRD2 upregulation by JQ1 across additional cancer types. **(A)** Western blot analysis of BRD2 in brain and breast cancer cells treated with DMSO or JQ1. **(B–C)** RT-qPCR and immunoblot analysis of BRD2 expression in human and mouse blood cancer cells, showing increased BRD2 levels upon JQ1 treatment (1 µM, 24 hours).

**Supplementary Fig. 2.**
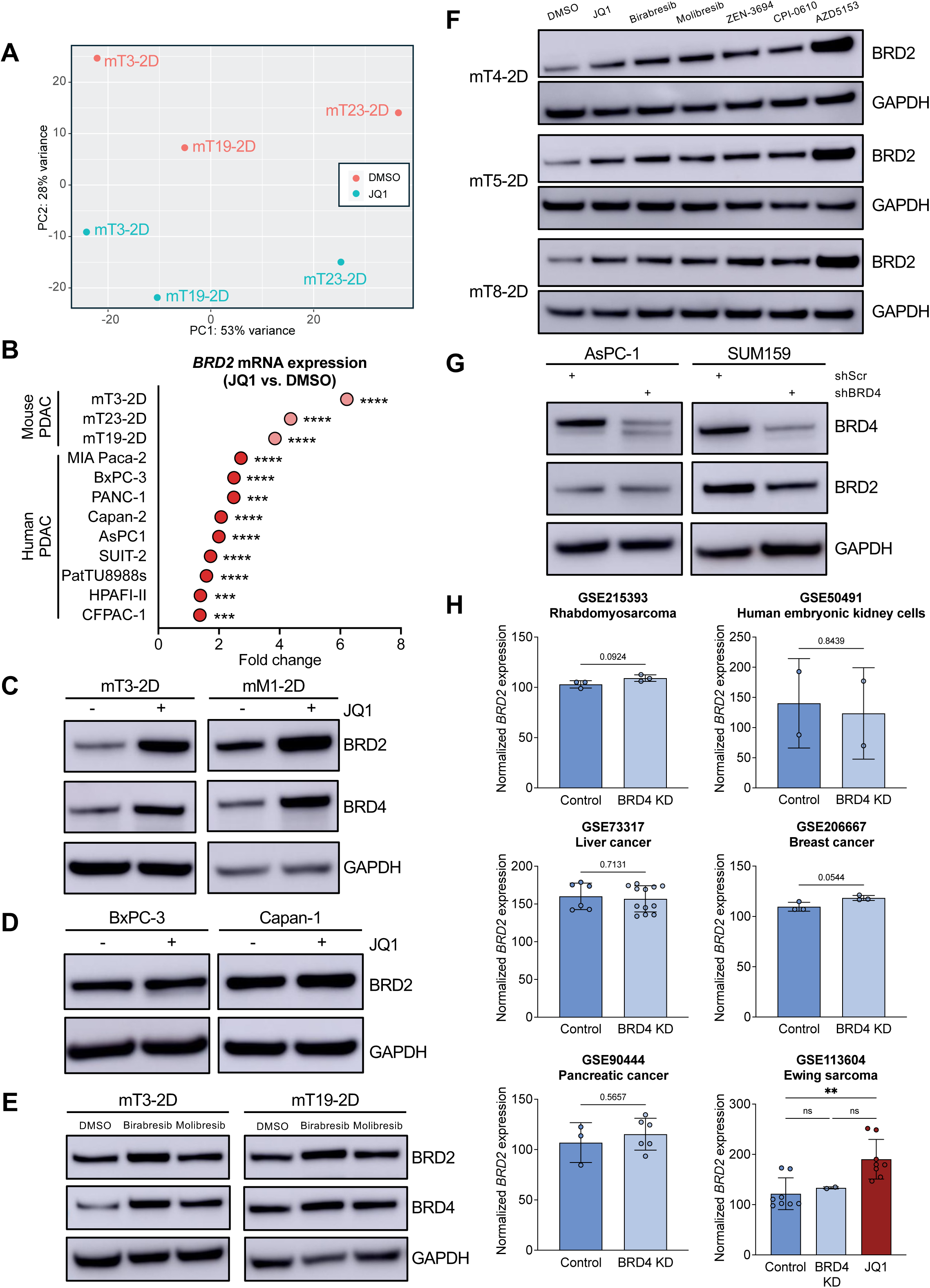
BRD2 induction by BET inhibition but not by BRD4 KD. **(A)** PCA of RNA-seq profiles from three different murine PDAC cell lines treated with DMSO or JQ1, showing distinct clustering of JQ1-treated samples. **(B)** RT-qPCR analysis of *BRD2* expression in multiple murine and human PDAC cell lines treated with JQ1, revealing significant induction across models. Statistical significance was determined using a two-tailed Student’s t-test (***p < 0.001, ****p < 0.0001) **(C–D)** Western blot analysis of BRD2 and BRD4 expression in murine (mT3-2D, mM1-2D) and human (BxPC-3, Capan-1) PDAC cells after JQ1 treatment. **(E)** Immunoblot analysis of BRD2 and BRD4 expression following treatment with JQ1, Birabresib or Molibresib in mouse PDAC cells (mT3-2D and mT19-2D). **(F)** Western blot analysis of BRD2 expression in murine PDAC cell lines (mT4-2D, mT5-2D, and mT8-2D) treated with different BETi (JQ1, Birabresib, Molibresib, ZEN-3694, CPI-0610, AZD5153), showing consistent BRD2 upregulation across inhibitors compared to control. **(G)** Western blot analysis of BRD2 expression after shRNA-mediated BRD4 KD in PDAC AsPC-1 and breast cancer SUM159 cells, showing that BRD4 depletion did not result in BRD2 induction by JQ1. **(H)** Analysis of publicly available RNA-seq datasets (GEO) across multiple cancer types, showing that BRD4 KD generally did not increase *BRD2* expression, whereas JQ1 significantly induced *BRD2*. Statistical significance was determined by two-tailed Student’s t-test (**p < 0.01).

**Supplementary Fig. 3.**
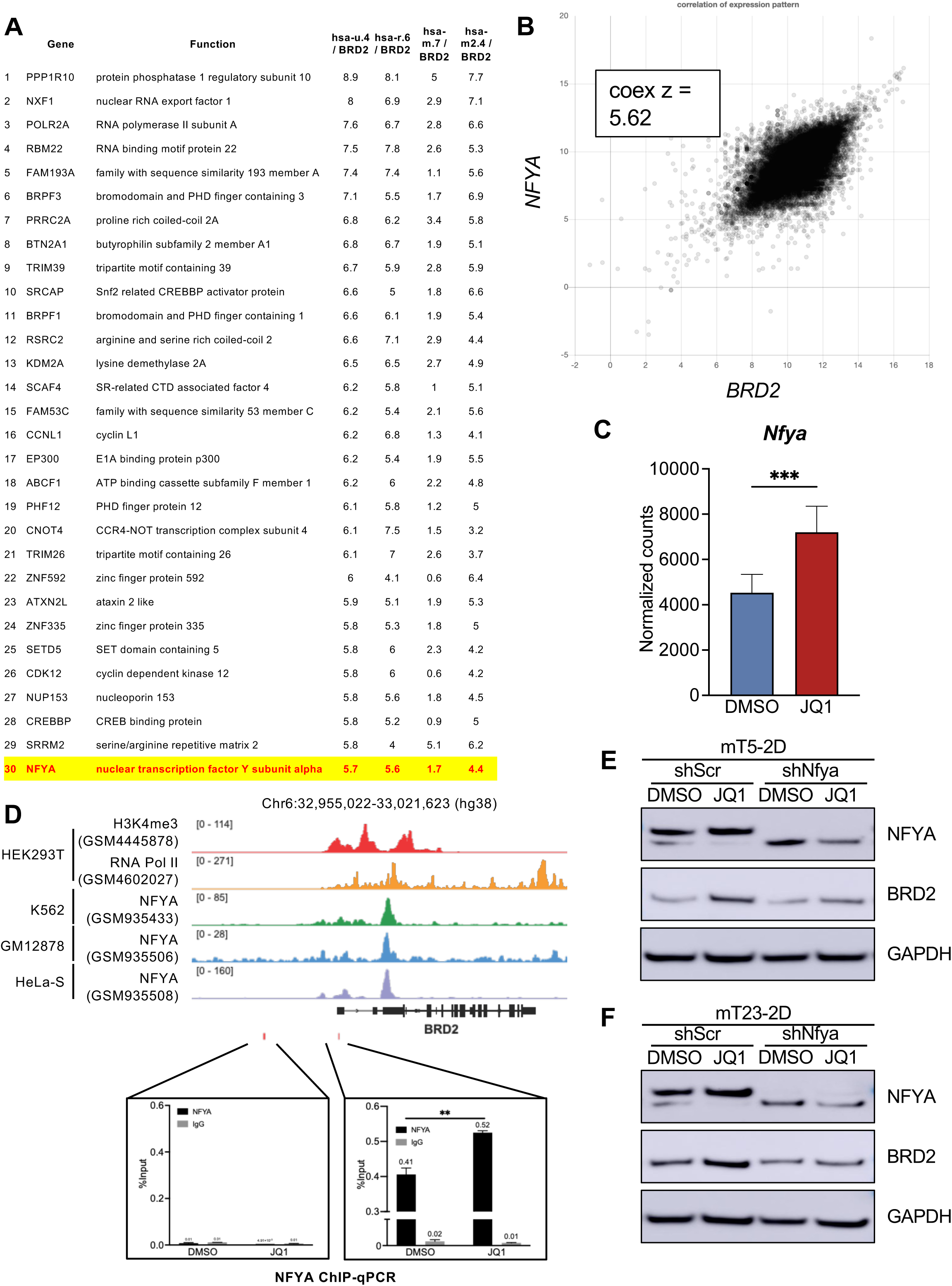
NFYA emerges as a candidate regulator of BRD2. **(A)** List of top 30 BRD2-interacting transcriptional regulators identified from COEXPRESdb. **(B)** Co-expression analysis of NFYA and BRD2 using COEXPRESdb, showing a strong positive correlation across transcriptomes (z = 5.62). **(C)** RNA-seq analysis of *Nfya* expression in mouse PDAC cells treated with DMSO or JQ1, showing significant *Nfya* upregulation upon JQ1 treatment. Statistical significance was determined by two-tailed Student’s t-test (***p < 0.001). **(D)** Public ChIP-seq tracks showing H3K4me3, RNA pol II, and NFYA binding across multiple cell types at the BRD2 locus. Lower panels: NFYA ChIP-qPCR in HEK293T cells under DMSO or JQ1 treatment showing enrichment at the *BRD2* exon 1 region relative to IgG and an intergenic negative control. Statistical significance was determined by two-tailed Student’s t-test (**p < 0.01). **(E–F)** Western blot analyses showing NFYA and BRD2 expression in control (shScr) and NFYA-KD (shNfya) murine PDAC lines mT5-2D (E) and mT23-2D (F) following DMSO or JQ1 treatment. NFYA depletion reduces BRD2 induction upon BET inhibition, which is consistent with human cell line data.

**Supplementary Fig. 4.**
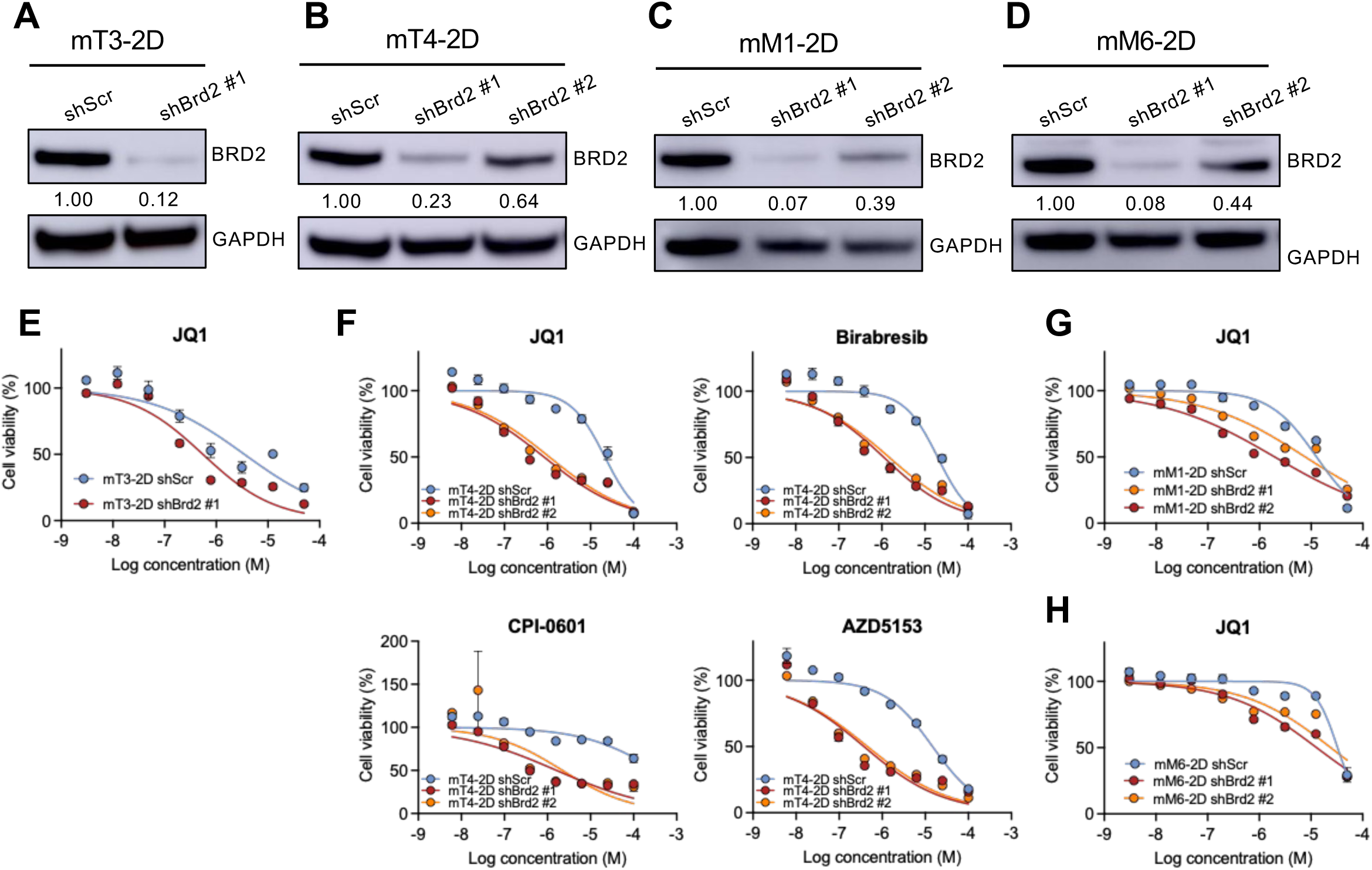
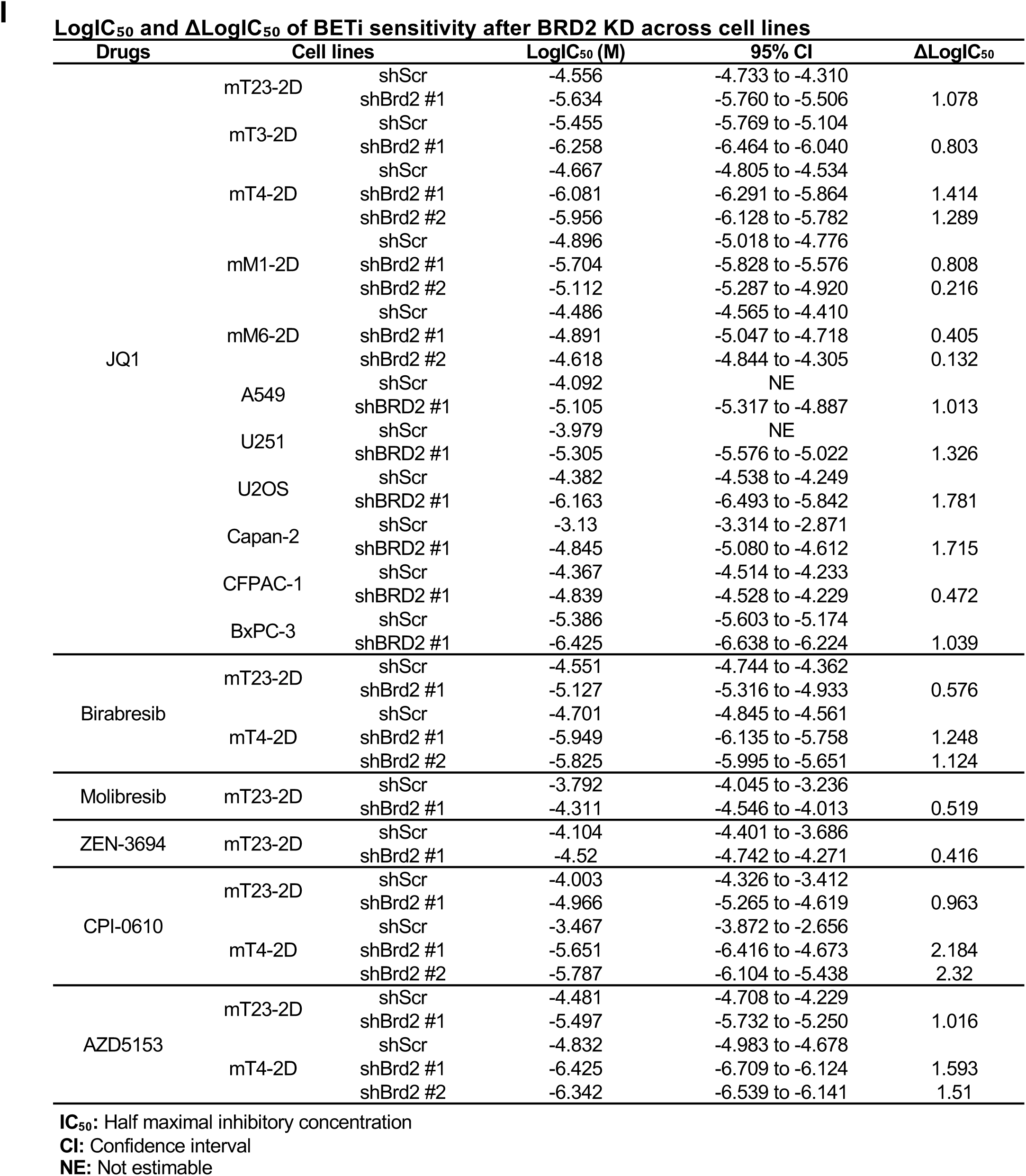
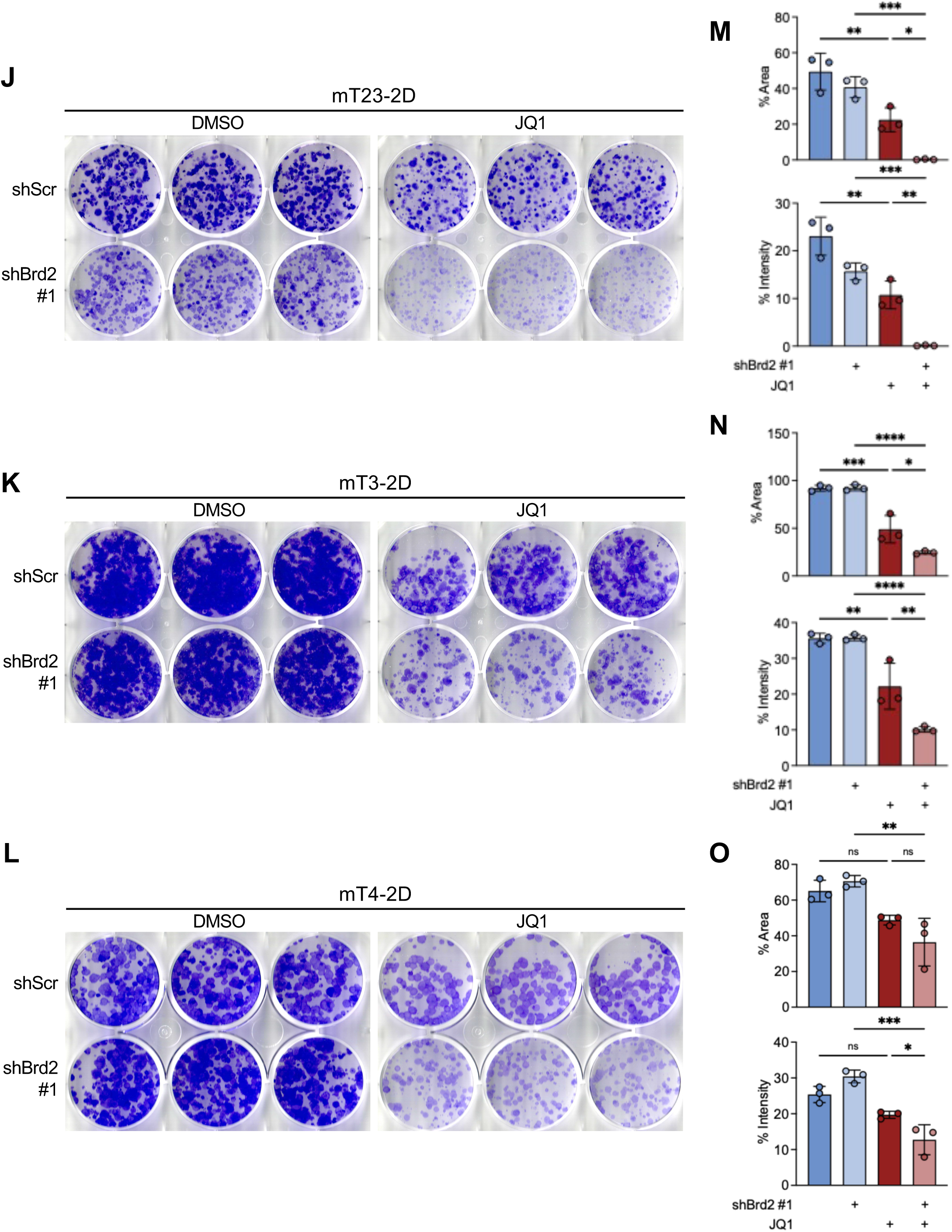
BRD2 KD enhances sensitivity to BET inhibition across murine PDAC models. **(A–H)** Immunoblots validating BRD2 KD using independent shRNA constructs (shBrd2 #1 and/or shBrd2 #2, as indicated) and dose–response curves for BETi showing reduced cell viability in BRD2-depleted cells compared with shScr controls in mouse PDAC cell lines. **(I)** Table of dose–response analyses showing LogIC_50_ values with 95% confidence intervals (CI) and ΔLogIC_50_ values for BETi sensitivity following BRD2 KD across cell lines (ΔLogIC_50_ calculated relative to shScr within each cell line). **(J–O)** Representative colony formation assays and their quantification (colony area and staining intensity) in mouse PDAC cell lines.

**Supplementary Fig. 5.**
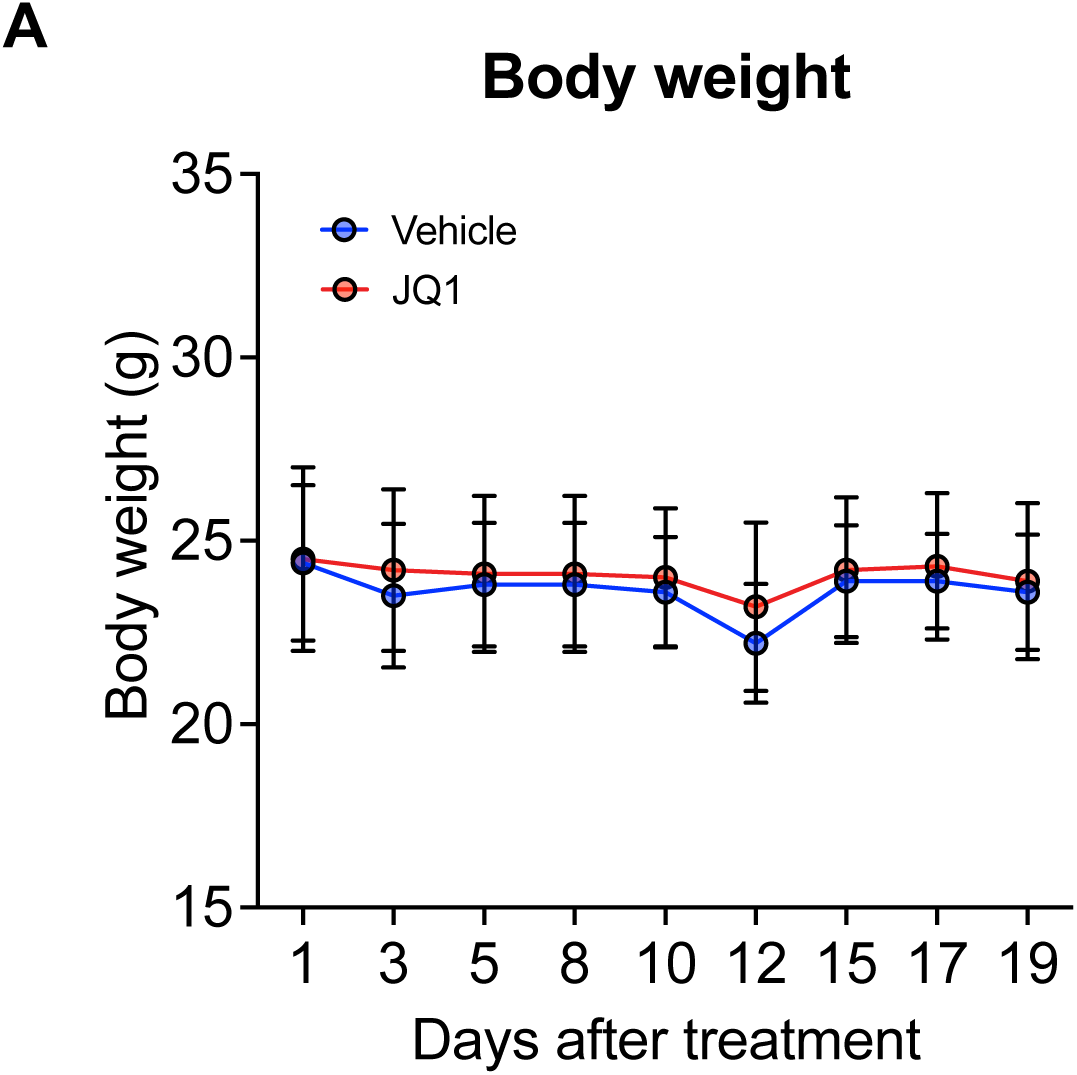
Body weight of mice was maintained during JQ1 treatment. **(A)** No significant weight loss was observed in mice bearing tumors treated with vehicle or JQ1.

**Supplementary Fig. 6.**
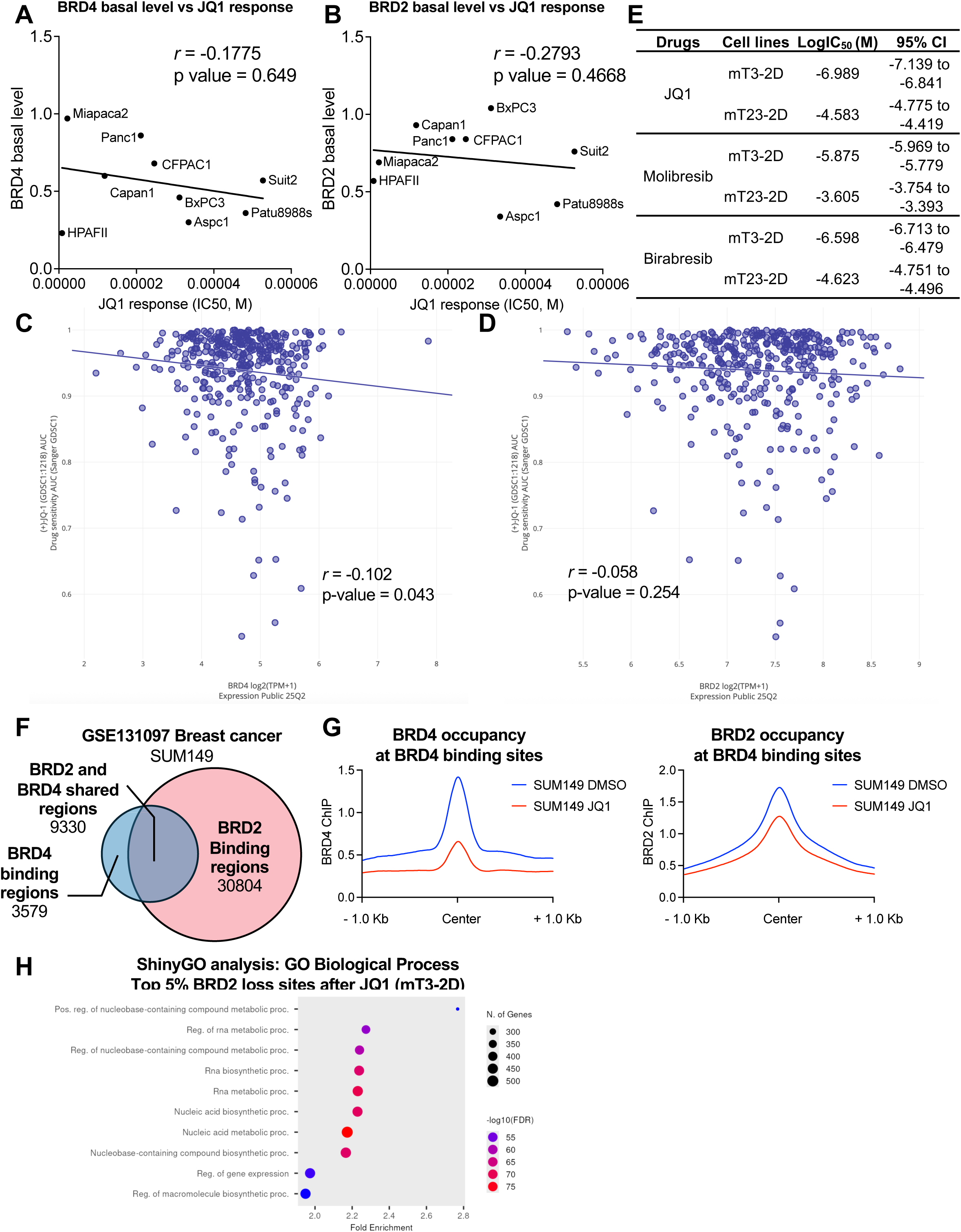
Basal BRD4/BRD2 levels and chromatin occupancy do not predict JQ1 response. **(A–B)** Correlation analyses between basal BRD4 (A) or BRD2 (B) protein expression levels and JQ1 sensitivity across human PDAC cell lines. No significant associations were observed. **(C–D)** Correlation of *BRD4* and *BRD2* mRNA expression levels with JQ1 sensitivity across public cancer transcriptome datasets. Both factors showed weak or no predictive value. **(E)** Summary table of LogIC_50_ and 95% CI for JQ1, Molibresib, and Birabresib in BETi-sensitive (mT3-2D) and BETi-resistant (mT23-2D) cells. **(F)** Venn diagram of BRD2 and BRD4 ChIP-seq binding regions in TNBC (GSE131097, SUM149), highlighting substantial overlap in line with our PDAC findings. **(G)** Metaprofiles of BRD4 (left) and BRD2 (right) occupancy at BRD4-bound sites under DMSO or JQ1 conditions, suggesting BRD2 binding persists at regions where BRD4 occupancy is displaced upon JQ1. **(H)** GOBP enrichment analysis (ShinyGO) of the top 1,000 JQ1-responsive BRD2-bound regions annotated by GREAT, revealing significant enrichment for RNA metabolic and biosynthetic processes.

## Conclusions

Our study uncovered BRD2 upregulation as a conserved adaptive resistance mechanism to BET inhibition across diverse cancer types. Through integrated transcriptomic and epigenomic analyses, we demonstrated that BETi treatment induced BRD2 expression at the transcriptional level, at least in part mediated by the NFYA transcription factor. BETi-mediated BRD2 induction acts as a compensatory mechanism, diminishing the antitumor efficacy of BETi. Notably, BRD2 KD sensitizes cancer cells to BETi, suggesting the therapeutic potential of dual targeting strategies that inhibit BRD2 alongside BETi to improve patient outcomes across multiple cancer types.

## Declarations

### Ethics approval and consent to participate

All animal studies were conducted in accordance with institutional and national guidelines and approved by the University of California, Davis (UC Davis) Institutional Animal Care and Use Committee (IACUC; Protocol #24016, approved August 22, 2024). UC Davis is accredited by the Association for Assessment and Accreditation of Laboratory Animal Care International (AAALAC) and maintains an Animal Welfare Assurance with the Office of Laboratory Animal Welfare (OLAW; Assurance Number D16-00272, A3433-01).

### Consent for publication

All authors have reviewed and approved this manuscript and consent to its publication.

### Availability of data and material

Raw and processed RNA-seq and ChIP-seq data generated in this study are available in GEO under accession numbers GSE307284 (RNA-seq) and GSE307285 (ChIP-seq).

### Authors’ contributions

S.A. and C.I.H. conceptualized and designed the study. S.A., J.J., and E.L. performed the experiments and analyzed the data. S.A. drafted the original manuscript, and C.I.H. reviewed and edited the manuscript. All authors read and approved the final version of the manuscript.

## Abbreviations

AML: acute myeloid leukemia
AT/RT: atypical teratoid/rhabdoid tumor
BC: breast cancer
BET: bromodomain and extra-terminal domain
BETi: BET inhibitors
BL: Burkitt lymphoma
CC: cervical cancer
CML: chronic myeloid leukemia
CRC: colorectal cancer
CRPC: castration-resistant prostate cancer
ES: Ewing sarcoma
ET: extra-terminal
GBM: glioblastoma
GIST: gastrointestinal stromal tumor
HCC: hepatocellular carcinoma
HNSCC: head and neck squamous cell carcinoma
KD: knockdown
Lum A: luminal A
MEL: melanoma
MM: multiple myeloma
NB: neuroblastoma
NFYA: Nuclear Transcription Factor Y subunit Alpha
NSCLC: non-small cell lung cancer
OC: ovarian cancer
PDAC: pancreatic ductal adenocarcinoma
RMS: rhabdomyosarcoma
SCLC: small cell lung cancer
SE: super-enhancers
TAD: topologically associating domain
T-ALL: T-cell acute lymphoblastic leukemia
TF: transcription factors
TNBC: triple-negative breast cancer

## Acknowledgements

This study was supported by National Cancer Institute grant 5R37CA249007 (C.-I.H.). The authors thank all members of the Hwang lab for their support throughout the course of this study, particularly Jihao Xu and Minh Duc Pham for their valuable discussions. The authors are also grateful to Dr. Chayarndorn Phumsatitpong for assistance with imaging the tumor and to Kedi Huang for help in optimizing the R code for generating the 3D volcano plot. S.A. was supported by the Anandamahidol Foundation (Thailand) and University of California Davis Comprehensive Cancer Center Director’s Fellowship. J.J. was supported by Provost’s Undergraduate Fellowship.

## Authors’ information

### Authors and affiliations

Department of Microbiology and Molecular Genetics, College of Biological Sciences, University of California, Davis, Davis, CA, United States

University of California Davis Comprehensive Cancer Center, Sacramento, CA, United States

### Corresponding author

Correspondence to Chang-il Hwang

